# The microprotein SEP^53BP1^: its bizarre mode of translational expression and intracellular behaviour

**DOI:** 10.64898/2026.05.04.722586

**Authors:** Joseph A. Curran, Killian A.J. Curran, Marta A. Inchingolo, Pascale Jaquier-Gubler

## Abstract

Microproteins are proteins of <100 amino acids. They represent a major, and until recently, overlooked fraction of the human proteome. However, it has now been demonstrated that many of these proteins play key roles in cellular physiology. Our group reported the expression of a microprotein expressed from an ioORF within the 53BP1 CDS arising as a result of delayed translational reinitiation mediated by a small uORF within the 5’ TL. We named this microprotein SEP^53BP1^. We have sought to expand these studies with the ultimate aim of establishing a function for this microprotein. Although this remains elusive, we report findings providing new insights into the elements regulating its translation and demonstrate that the SEP^53BP1^ sequence serves as a Golgi targeting tag. Lastly, despite the fact that subunits of the proteasome feature prominently on interactome studies we were unable to demonstrate an impact of microprotein over-expression on the activities of both the proteasome and immunoproteasome.

## INTRODUCTION

The potential biological importance of the proteomic “dark matter” or “Ghost proteome’ is only beginning to be elucidated (1,2). These are a highly abundant group of proteins of less than 100 amino acids (aas), frequently referred to as microproteins, that have, because of technical reasons, been overlooked in classical mass-spectrometry based proteomic analysis. However, things have undergone a revolution largely because of ribosomal profiling (3–8). This technique revealed the extensive cellular microprotein proteome arising from the translation of small open reading frames (smORFs) within the mRNAs 5’ untranslated region (UTR, also referred to as transcript leader-TL), 3’ UTR, internal and overlapping ORF(s) embedded within the main CDS (ioORFs) but also many long noncoding RNAs (IncRNAs). These microproteins are also referred to in the literature as SEPs (small ORF encoded proteins), alternative open reading frames (Alt-ORFs), AltProts or micropeptides (9). Microproteins are notoriously difficult to study (6,10). Their small size means that they rarely present any well-defined protein domain, which complicates functional studies. They also tend to be relatively unstable and individually poorly abundant.

The mRNA 5’ TL is a key element in the regulation of the translational output at both the quantitative (total protein yield per mRNA) and qualitative (the nature of the proteins expressed) levels (11). Heterogeneity within the 5’ TL arises from alternative promoters and/or alternative splicing within the 5’ TL, impact on the translational readout and, as a consequence, the cellular phenotype (11,12). Furthermore, 5’ TL complexity is significantly amplified by the fact that most mammalian promoters initiate not from a single TSS, but rather a cluster of start sites within the core promoter. This is more marked in CpG (TATA-less) promoters that can exhibit TSS sites spanning XX nts (13–15). Complexity within the transcript population is further filtered by the ribosome during mRNA recruitment, the ribosome filter hypothesis (16), to fine-tune the proteome (17).

In a comparative transcriptome/translatome analysis performed on the tumoural MCF7 and non-tumoural MCF10A cell lines our lab identified over 550 genes exhibiting differential translation efficiency (TE) (18). Within this group, we observed distinct differential promoter usage patterns in both the transcriptome and translatome, behaviour consistent with the ribosome filter hypothesis (16,19,20). One gene that drew are attention was *TP53BP1*, whose main CDS codes for a protein, 53BP1, central in DNA repair (21). *TP53BP1* uses two promoters (P1 and P2) that give rise to alternative 5’ TL transcript variants. P1 generates transcripts V1/2 (NM_001141979.1, NM_001141980.1), that differ by alternative splicing within the ORF. These are generally the major transcripts in most cell types. The 5’TL is 113nts long, 71% G/C and contains no uAUGs. We obseved V1/2 on the polysomes of both MCF7 and MCF10A cells (18). The second TL, V3 (NM_005657.2), arises from P2. Based on both the CAGE and EPD databases, it has a 5’TL of 278nts, is 62% G/C and carries a small 5 codon uORF 15 nts upstream of the 53BP1 start codon (AUG^53BP1^). V3 was polysomal in the MCF7 but non-polysomal in MCF10A cells, pointing to a cell-type specific recruitment. We subsequently established that the small uORF in the V3 transcript permitted a robust delayed reinitiation event (22) at an ioORF overlapping the primary CDS (23). Curiously, the efficiency of this initiation event was coupled to transcript sequences upstream of the AUG start site. The product of this ioORF, a 50 aas protein that we named SEP^53BP1^, was detected in several cell lines including MCF7, THP1 and Raji (but not HeLa, HEK293T and MCF10A) and displayed both nuclear and cytoplasmic compartmentalisation. A fraction was also associated with internal cellular membranes (23). As with most microproteins, structural predictions revealed no clear functional domain. In the OpenProt database (https://openprot.org/), SEP^53BP1^ is referenced as AltORF (alternative ORF) #IP_232576, as peptides corresponding to this microprotein have been observed in mass spectrometry studies. Our earlier report finished with the establishment of an interactome for SEP^53BP1^. Both Y2H (yeast two-hybrid) and coIP-MS (co-immunoprecipitation mass spectrometry) approaches revealed that it associated with multiple cellular components including the protein turnover apparatus, in particular the exposed C-terminal of the α4 (PSMA4) subunit on the 20S proteasome barrel. Recent studies have indicated that proteins interacting with C-terminal of the α4 can facilitate the degradation of specific target substrates in both a ubiquitin dependent and independent manner (24,25). Could the proteasome-SEP^53BP1^ interaction modulate proteasome activity?

In this manuscript we have sought to complete our earlier work both by re-examining the curious role of the mRNA sequences upstream of the mRNA SEP^53BP1^ AUG start codon on its translational expression and, starting from the interactome data, seek to establish a cellular function for this microprotein. Although these objectives remain elusive, are studies have revealed a number of observations that shed some light on translational control and the behaviour of the microprotein sequence in living cells. It is to be hoped that these results may stimulate further investigations by other groups interested in microprotein expression and function.

## RESULTS/DISCUSSION

### Curiosities of SEP53BP1 translational expression

The SEP^53BP1^ microprotein arises from an ioORF within the 53BP1 CDS (18). One curiosity of its expression is the important requirement for the 53BP1 mRNA sequences upstream of the AUG^SEP^ start codon (sequences referred to as V3) (23). Microprotein expression was severely reduced when these sequences were removed and could not be restored by the substitution of a foreign sequence of similar length (the 3’ UTR of the ACOXL transcript) (23). This enigma remains troubling, even more so because the same V3 region was unable to promote the expression of synthetic smORFs of the same length as SEP53BP1, namely 50 codons or SEP^53BP1^ fusion proteins (see below). This might suggest some interplay between the *de-novo* synthesised microprotein and the V3 region. However, direct interactions between SEP^53BP1^ and V3 mRNA sequence remains to be explored. The “water is further troubled” by our new observation that the V3 sequence requirement can be mimicked by the presence of an upstream intron. This was initially observed when the SEP^53BP1^ ioORF, devoid of upstream and downstream 53BP1 sequences, was transferred from the “intron-less” CMV based expression vector pcDNA3 (in which most or our studies have been performed) to the pEBS-PL episomal Epstein-Barr virus-based mammalian expression vector which carries an SRα promoter and a downstream small intron upstream of the cloning site (26). (Figure 1A). To confirm that this was coupled to the intron rather than the promoter switch, we transferred the same intron into the pcDNA vector background in both orientations downstream of the promoter and upstream of the ioORF (without V3). Microprotein expression was observed when the intron was in the correct orientation and was not observed in the construct carrying the reversed intronic sequence (Figure 1B). It should be noted that the reverse intronic sequence did not introduce uAUG(s) that would act as translational repressors. However, expression did correlate with splicing of the intron as confirmed by RT-PCR on total cell RNA (Figure 1B). Intriguingly, the SEP^AUG^ start codon lies in the second exon of the mature transcript. This might suggest that elements of the splicing machinery that move into the cytoplasm with the mature transcript, or elements of the exon junction complex (EJC), are retained on the V3 sequence and interact with the translational machinery during microprotein expression or shortly after release (27–29). It has been observed that splicing enhances protein yield because the EJC deposited on the mRNA promotes polysomal association (28). However, how and why these elements are retained on the pcDNA3 V3-SEP transcript that carries no intron (only the mature EXON1-EXON2 V3 sequence) remains unclear.

**Figure 1:**
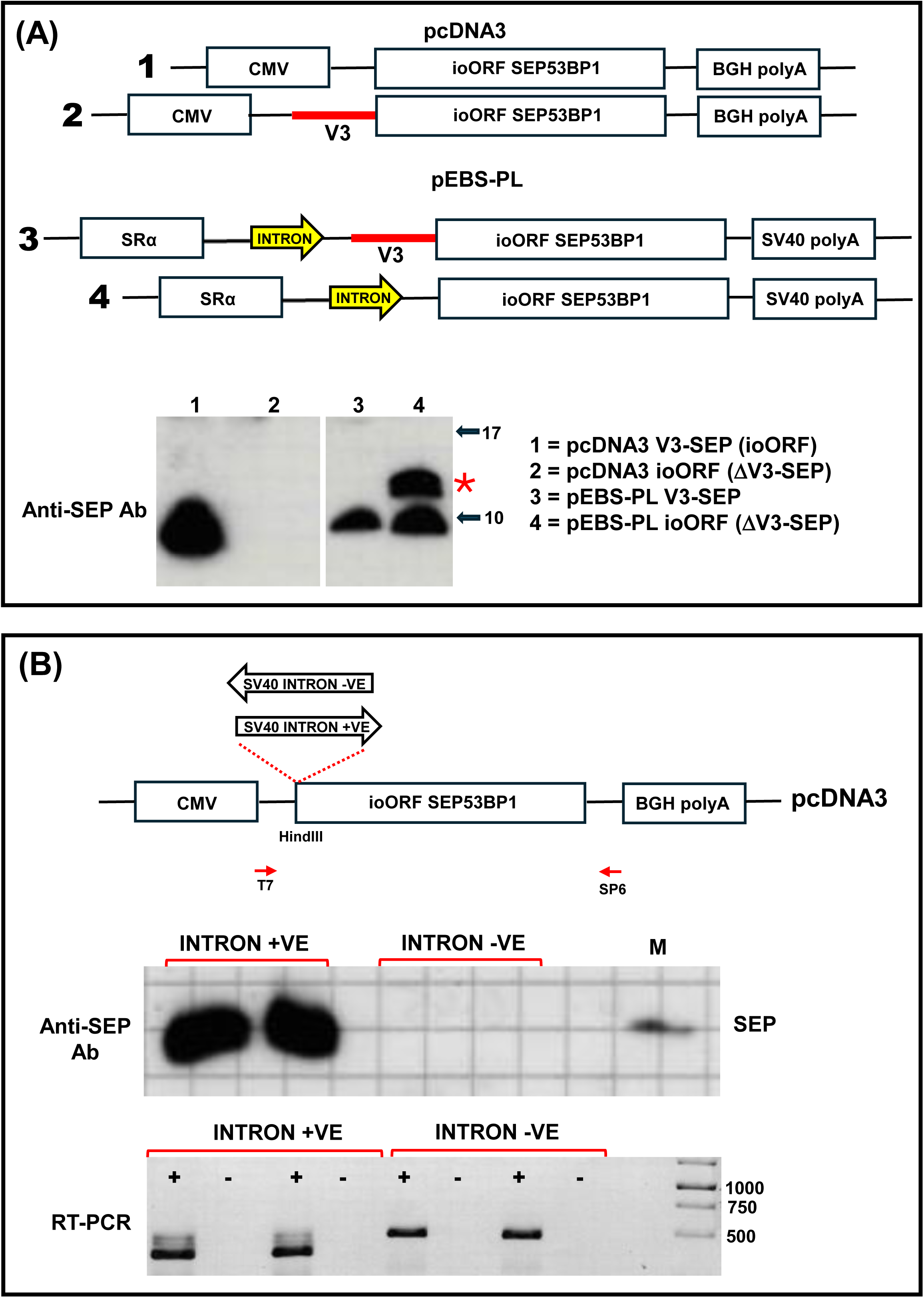
Introns promote the expression of the SEP^53BP1^ ioORF. (A). The upper panels are schematic images of the pEBS-PL and pcDNA3 ioORF expression vectors with or without the V3 upstream region. Indicated also is the small intron present in the pEBS-PL vector. The lower panel shows an immunoblot performed on cell extracts from transfected HEK293T cells using the anti-SEP antibody. The star indicates an additional band detected by the antibody whose origins remain obscure. (B). The SV40 intron from the pEBS-PL vector was transferred as a HindIII fragment into the pcDNA3 V3 minus background. This gave rise to intron insertions in both orientations (upper panel). SEP^53BP1^ expression in HEK293T transfected cells was detected by immunoblotting. Duplicate clones from both intron orientations were analysed. The marker M is the a synthetic SEP^53BP1^ peptide (middle panel). Total RNA was isolated from the transfected cell extracts and analysed by RT-PCR using the SP6 and T7 primers (- indicates the RT minus control). Products were analysed on an agarose gel (lower panel).

### Attempts to express synthetic microproteins

For functional studies with the SEP^53BP1^ protein we sought to generate a similar 50 aas synthetic microprotein as a negative control. This microprotein would carry epitope tags, allowing us to monitor its transient expression, and would carry an optimal Kozak context AUG initiation codon so as to assure maximal expression (30). We synthesised pcDNA3-based plasmid clones (BioCat GmbH and Gene Universal Inc) that could be employed to transiently express a series of synthetic microproteins (see Table 1), starting with a FLAG-2MYC construct (an N-terminal FLAG tag fused to two MYC tags: Table 1). However, on immunoblots using anti-FLAG and anti-MYC Abs we observed no protein expression (in these experiments the positive control was supplied by a FLAG/MYC construct, ΔKpn LP/SP) (31). We then introduced a stability sequence (STAB) (construct FLAG-MYC-STAB: Table 1) (32). Once again, we failed to observe microprotein (Supplemental Figure S1) despite robust intracellular mRNA expression in the transfected HEK293T cells as monitored by RT-PCR (Figure 2A). This was not improved by the introduction of a V3 upstream sequence or the introduction of either the N-terminal 15 aas or the C-terminal 14 aas of SEP^53BP1^ despite evidence of intracellular mRNA transcription in all cases (Figure 2A). Neither was a FLAG-MYC-STAB microprotein detected when translation was driven by the EMCV, HCV or CrPV IRESes (Figure 2B). This would suggest the proteins are highly unstable. Attempts to stabilise the microproteins in cells using the inhibitor MG132 failed, suggesting that if the protein is being rapidly degraded it is probably not via the proteasome (not shown). Mechanistically, what is blocking the expression of these synthetic microproteins remains unclear. Levels of intracellular mRNAs comparable to V3-SEP^53BP1^ (Figure 2A) and our inability to observe an effect with the proteasome inhibitor MG132 would tend to exclude nonsense mediated decay (NMD). This pathway serves to limit the expression of truncated proteins generated by mutations that introduce premature stop codons. NMD is generally triggered when translation terminates more than ∼50–55 nucleotides upstream of a post-splicing exon-exon junction during the pioneer round of translation The response induces rapid proteasome-mediated degradation of both the nascent polypeptide chain and the mRNA template (33–35). However, since our synthetic transcripts are derived from intron-less mRNAs that appear to be stable in the transfected cell, and protein expression is not observed upon proteasome inhibition, all appears to exclude this mechanism in the negative regulation of synthetic microprotein expression. Likewise, whereas HCV and EMCV IRES-mediated translation remains responsive to NMD this is not the case with the CrPV IRES, probably because the latter does not require the initiation factor eIF3 (36–38). Yet we still failed to observe microprotein expression when driven by the latter.

**Figure 2:**
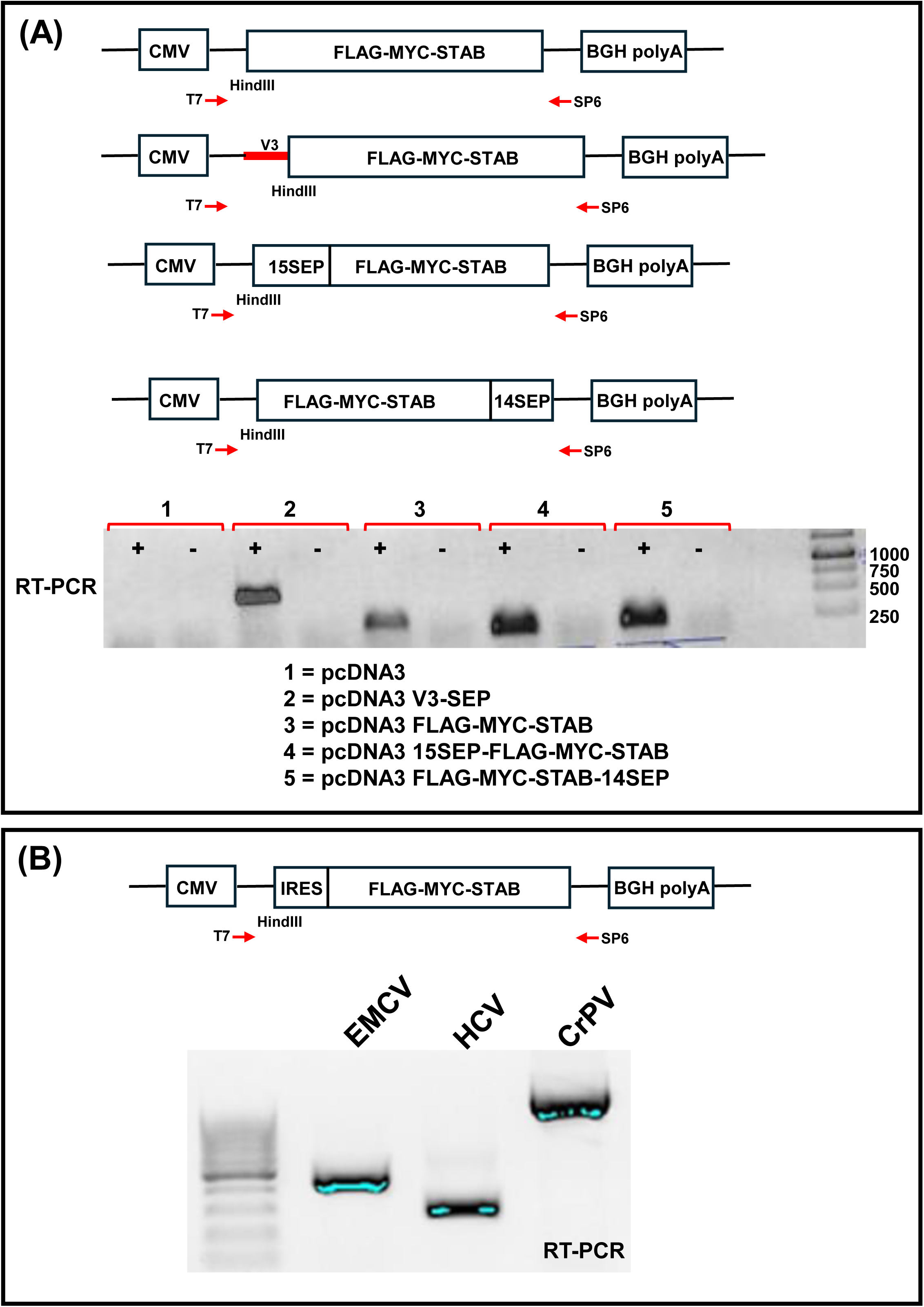
Attempts to express an epitope tagged 50 amino acid experimental control. (A) Initially we generated a 153 nts (50 codons plus STOP codon) FLAG-MY-STAB cassette that was cloned into a pcDNA3 vector with or without the V3 upstream sequence. This was subsequently modified with the introduction of the first 15 N-terminal codons of SEP^53BP1^ and the last 14 C-terminal codons of SEP^53BP1^ (upper panels). All constructs were transcriptionally expressed in transfected cells as confirmed by RT-PCR using the SP6 and T7 primers (- indicates the RT minus control: lower panel). However, we repeatably failed to detect protein expression on immunoblots with both the anti-FLAG and anti-MYC antibodies (see Supplemental Figure S1). (B). IRESes from EMCV (encephalomyocarditis virus), HCV (hepatitis A vurus) and CrPV (cricket paralysis virus) were inserted upstream of the SEP^53BP1^ AUG start codon (upper panel). Despite robust transcription in transfected cells (as determined RT-PCR using the SP6 and T7 primers: lower panel) no protein expression could be determined by immunoblotting.

**Table 1:**
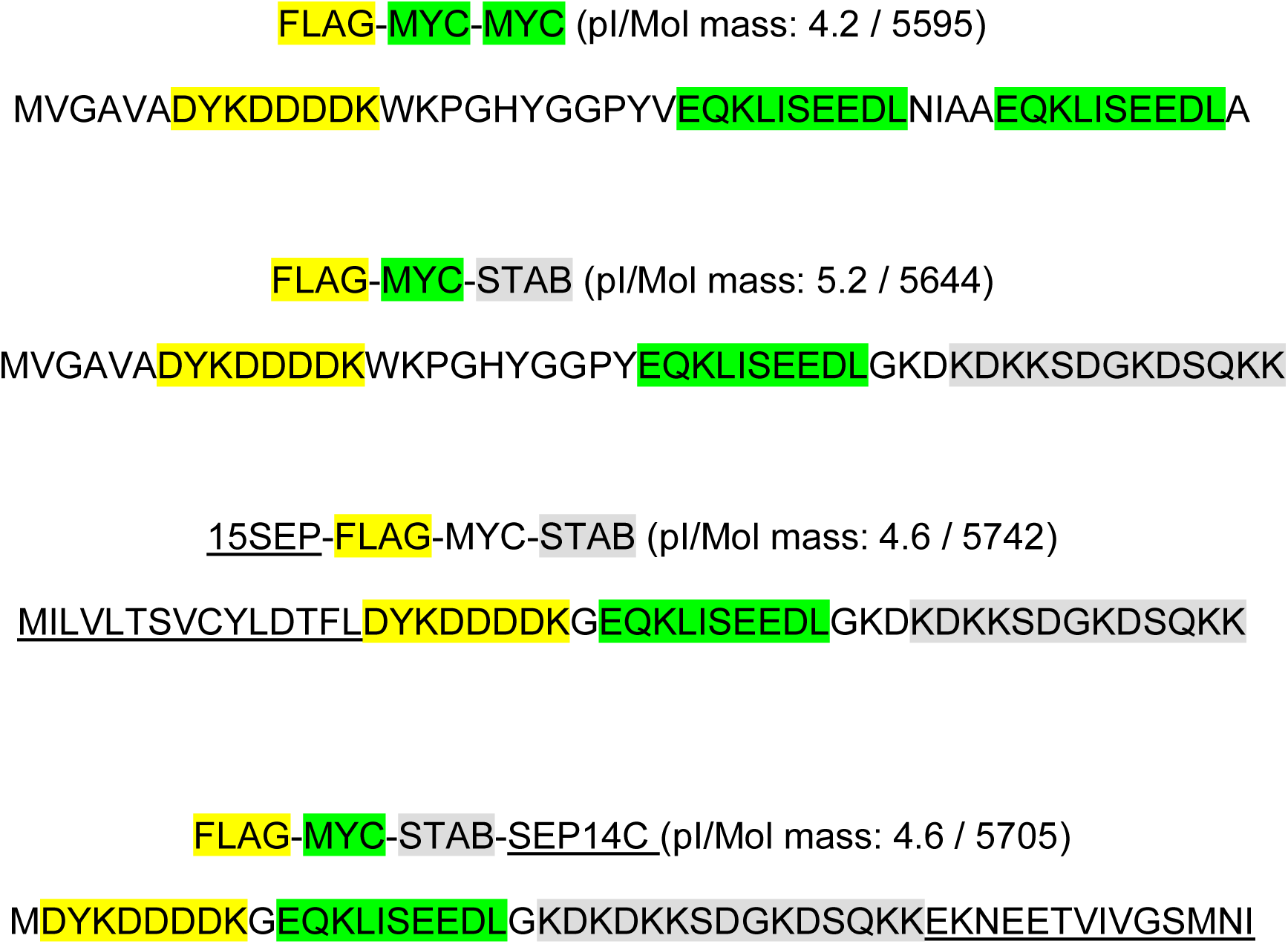
PRIMARY STRUCTURE SYNTHETIC MICROPROTEINS.

### Fusion proteins: The behaviour of the SEP53BP1 region

Microproteins are frequently studied by fusion to fluorescent tags, frequently GFP, despite the fact that these “tags” are considerably bigger than the microprotein itself (GFP is 238 aas), and can consequently alter microprotein behaviour (6). Fusion proteins can facilitate intracellular localisation studies, with the caveat that the GFP module is not always neutral in this regard (39). Nonetheless, we fused the SEP^53BP1^ sequence to the N-terminus (SEP-GFP) or C-terminus (GFP-SEP) of GFP in a pcDNA3 expression vector (Figure 3A). Transient expression in HEK293T cells gave robust transcription of all constructs as evidenced by RT-PCR (Figure 3B). Protein expression was monitored by immunoblotting using initially a commercial anti-GFP monoclonal Ab (Roche: GFP#1) and our anti-SEP polyclonal Ab (Figure 3B). The GFP#1 Ab detected expression of GFP and SEP-GFP but not GFP-SEP and, curiously, the SEP-GFP band co-migrated with GFP (Figure 3B). Things got even more bizarre when the same extracts were analysed with the anti-SEP polyclonal Ab. On this immunoblot, we observed a band only in the GFP-SEP transfections, migrating slower than GFP, consistent with the presence of the 50 amino acid SEP^53BP1^ sequence. However, if we loaded more protein extract onto the immunoblot, we frequently observed a weak ladder of bands in the SEP-GFP cells spanning the GFP-SEP to GFP molecular mass range (Figure 3B). This could suggest N-terminal trimming of the SEP sequence (but see below for an alternative explanation). We then repeated the analyse with two alternative anti-GFP Abs. Ab GFP#2 was a second commercial monoclonal (Abcam) whereas Ab GFP#3 (Proteintech) was a commercial rabbit polyclonal raised against a GFP fusion protein. The Ab GFP#2 detected expression only in the GFP transfected cells, whereas Ab GFP#3 detected expression in all extracts, including GFP-SEP (Figure 3C). SEP-GFP and GFP-SEP produced bands on the immunoblot of different size even though their amino acid composition and molecular mass are in principle identical (Mol.mass 33240; pI 6).

**Figure 3:**
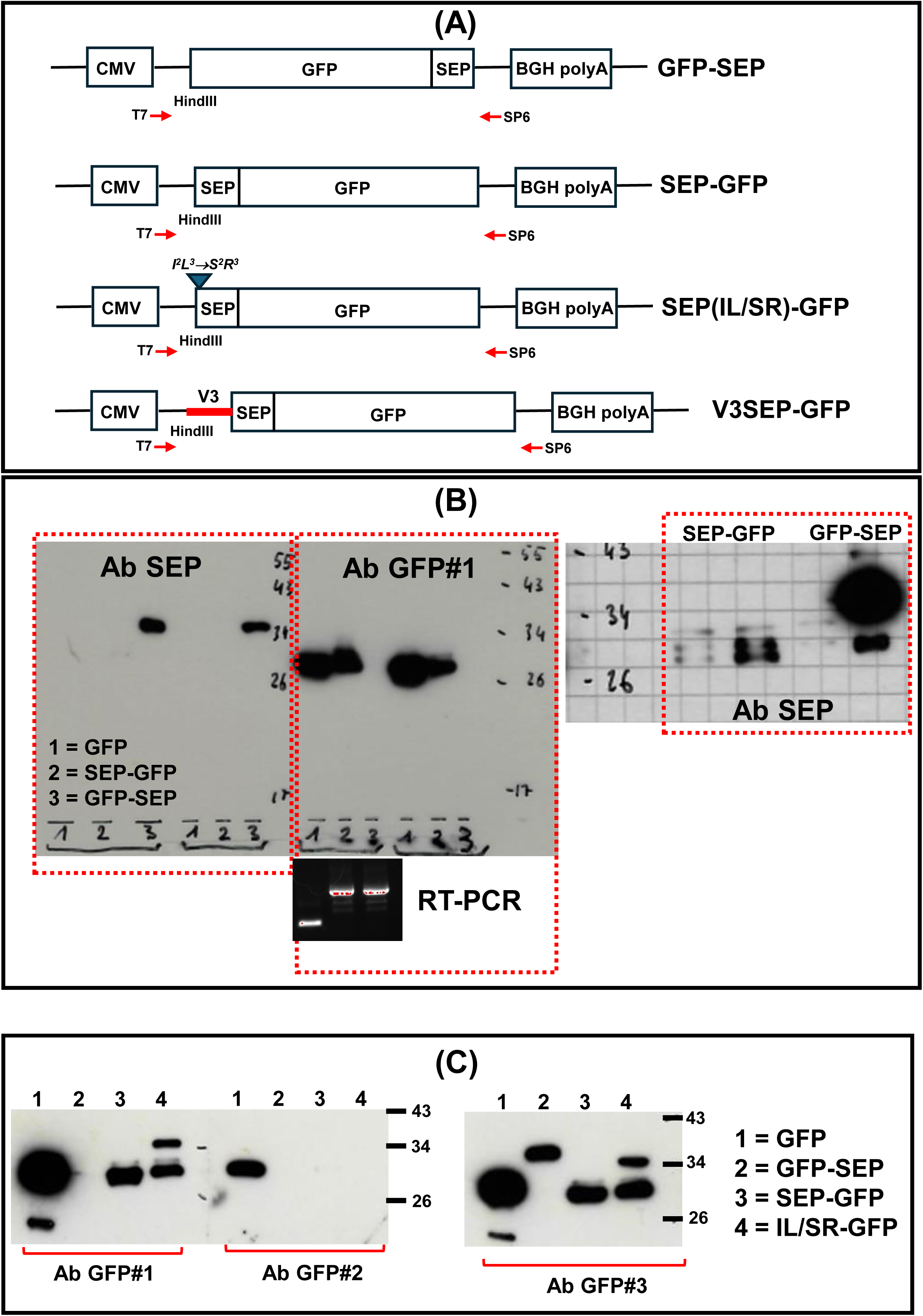
The behaviour of SEP^53BP1^-GFP protein fusions. (A). Schematic representation of the various protein fusions that were generated in a pcDNA3 plasmid background. (B) Constructs were expressed in HEK293T cells and protein expression monitored by immunoblotting with the anti-SEP polyclonal Ab and an anti-GFP monoclonal Ab (referred to as Ab GFP#1. left-hand columns). Intracellular transcription from all vectors was robust as evidenced by RT-PCR analysis using the T7 and SP6 primer set (lower panel insert). SEP-GFP and GFP-SEP extracts were re-analysed by immunoblotting with the anti-SEP Ab but with a greater protein load (40 μg) and a longer immunoblot exposure time (right-hand column). (C) The cell extracts were analysed for GFP expression using two additional anti-GFP Abs, namely, a second monoclonal (GFP#2) and an anti-GFP polyclonal (GFP#3). Included in this blot is an N-terminal mutant in the SEP-GFP construct referred to as IL/SR-GFP.

SEP-GFP protein expression levels were not enhanced by the introduction of the upstream V3 region (Figure 4A); therefore, the effect of this region with regards to translational expression is limited to the isolated ioORF. The low-resolution intracellular staining pattern of SEP-GFP mirrored that of GFP (Figure 4B), with the lower intensity probably reflecting the level of transcript (Figure 4A: the AUG initiation codon in each construct is optimal). Fluorescence was observed throughout the cell. However, the picture with the GFP-SEP was somewhat surprising. Despite robust protein expression, as demonstrated in immunoblots using the Ab GFP#3 (Figure 4B), fluorescence was barely detectable at camera settings that gave saturating signals with the other constructs (Figure 4B). The weak signal was also very punctate, reminiscent of Golgi staining. Using live cell imaging with an mCherry marker for the Golgi (GalT-mCherry, GalT encoding the amino acids 1-81 from β1,4-galactosyltransferase) we observed significant co-localisation (Figure 4C). Higher resolution images analysed using the Imaris software confirmed a considerable overlap between the GFP and mCherry signals (Figure 4D.1) with image deconvolution indicating that a fraction of SEP-GFP was localising to the exterior surface of the Golgi (Figure 4D.2). This live cell staining pattern and co-localisation was not observed in cells expressing GFP or SEP-GFP, whose behaviour were very similar i.e. diffuse GFP staining throughout the cell (Supplemental Figure S2).

**Figure 4:**
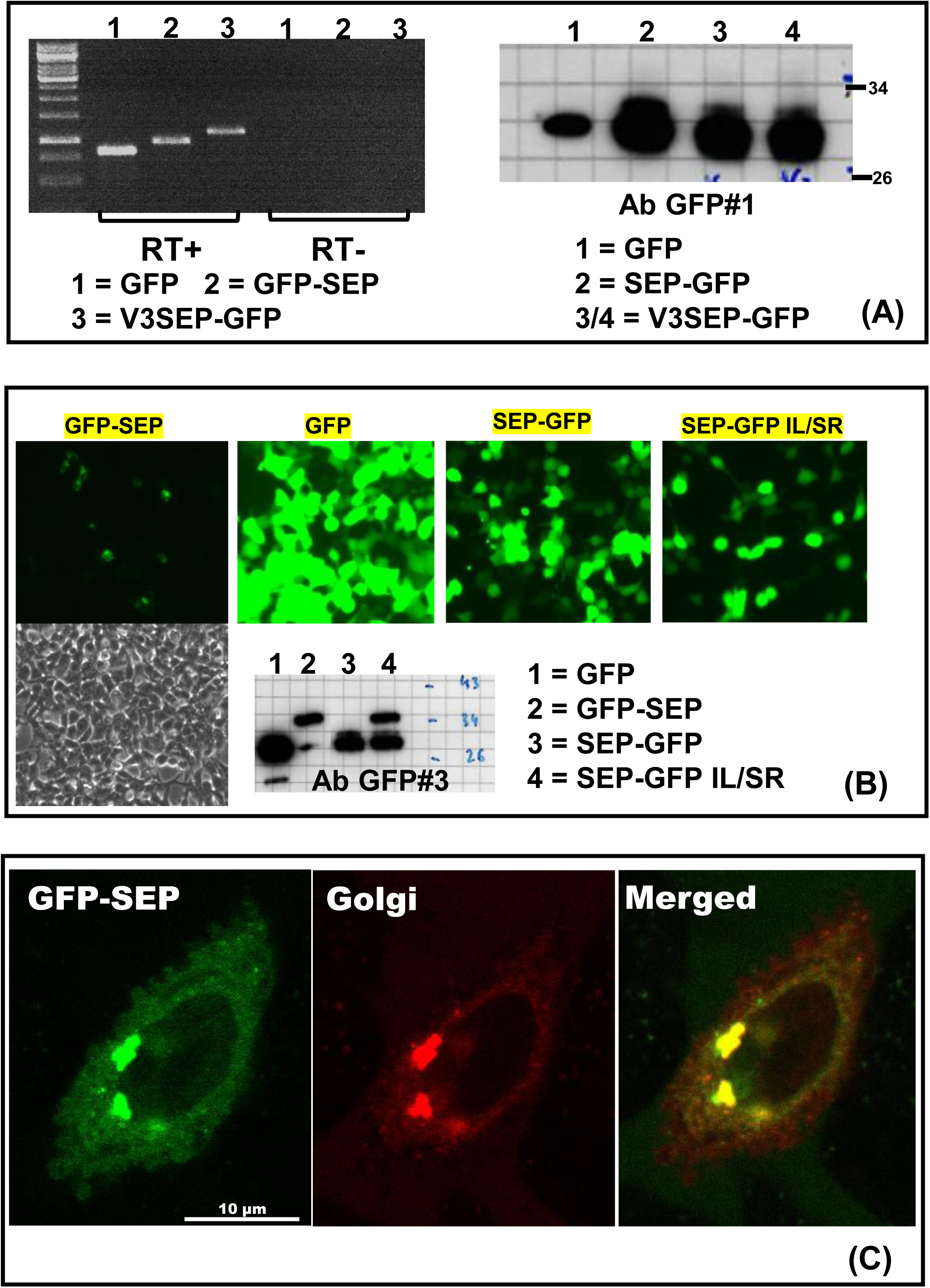
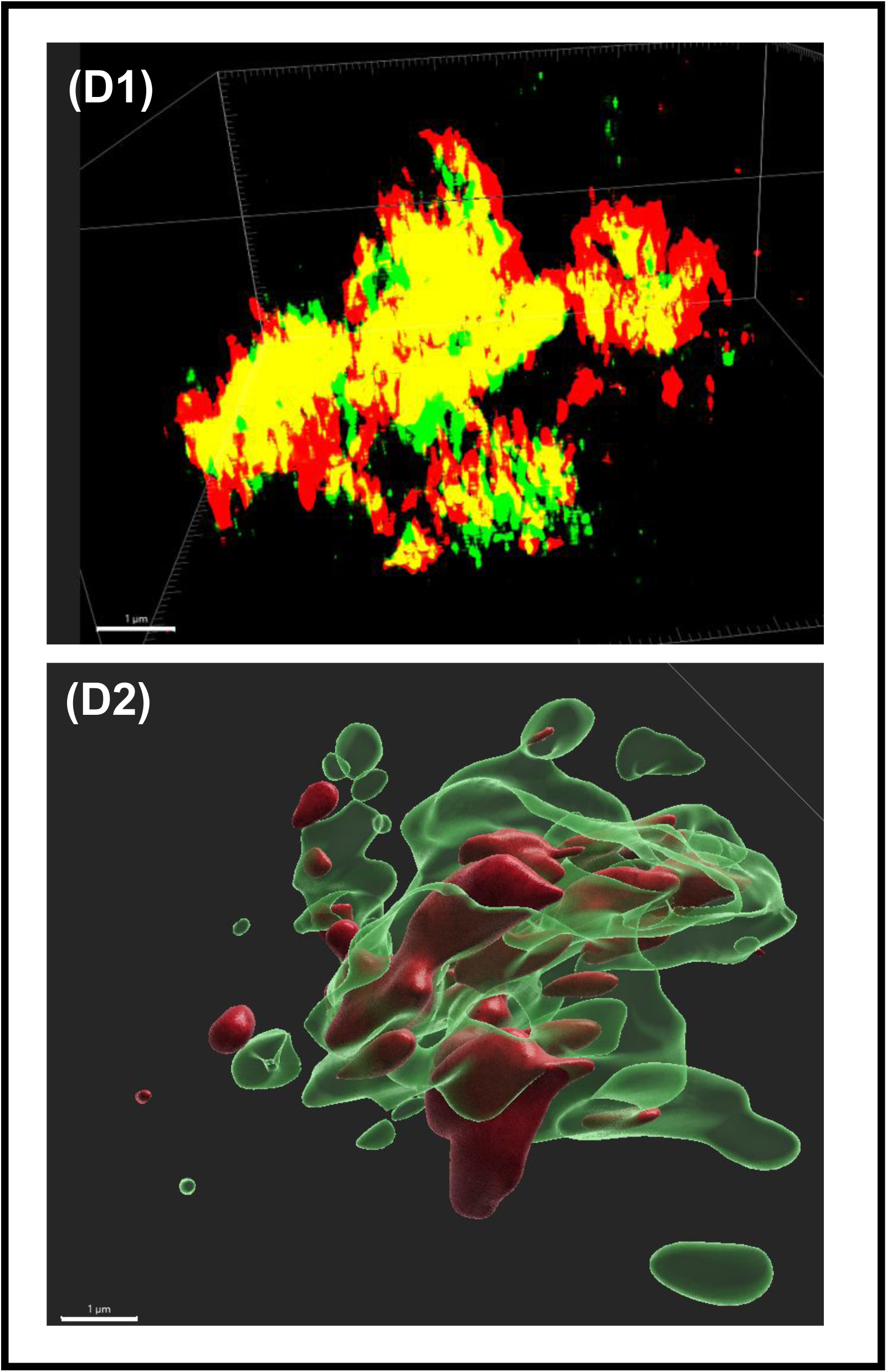
Intracellular behaviour of the GFP-SEP fusion constructs in HEK293T transfected cells. (A). On the left-hand side is an RT-PCR analysis of transcript expression performed on total cell RNA. Included is an RT minus (RT-) control. On the right is an immunoblot performed with the polyclonal anti- GFP#3 Ab. (B). Low magnification GFP fluorescence in transfected cells. Camera settings were the same for all the images. A live cell image is shown for the GFP-SEP transfected cells. The insert is an immunoblot of the transfected cells performed with the GFP#3 Ab. (C). Images taken from cells co-transfected with GFP-SEP and pmCherry-N1-GalT (expressing the Golgi marker β1,4-galactosyltransferase: Addgene). The merged image is indicated on the right. (D). High magnification live cell images focusing on the Golgi. The D1 panel depicts the raw image with the Golgi marker indicated in red, the GFP-SEP signal in green and the overlap in yellow. In the D2 panel, the Golgi image has been deconvoluted using the Imaris software to highlight the presence of the GFP-SEP protein on the outer surface of the Golgi.

The different migration of the SEP-GFP and GFP-SEP proteins and the very different behaviour of each in transfected cells was intriguing. The co-migration of SEP-GFP with GFP, the fact that SEP-GFP was not detected with the anti-SEP Ab and their very similar intracellular fluorescent profiles, suggested that a major part, or maybe all, of the SEP^53BP1^ region was missing on the SEP-GFP protein. This could arise translationally (leaky scanning downstream of the AUG^SEP53BP1^: this site is indicated as M^1^ in Figure 5A) or post-translationally (proteolytic processing of the SEP-GFP protein). Mutating the GFP initiation codon AUG^GFP^ (M^GFP^) to GCG (an Ala codon: M^GFP^/A) did not alter protein expression from the SEP-GFP construct (Figure 5A, lane 4), indicating that the band we observe in this construct that co-migrates with GFP does not arise from initiation events at this site. A similar mutational change at the AUG^SEP^ start site (M^1^/A: Figure 5 lane 5) or even the double mutant (M^1^/A M^GFP^/A: Figure 5 lane 6) did not alter expression. However, there are two additional start sites in the SEP-GFP ORF (M^27^ and M^48^) that are probably employed as initiation codons when the AUG^SEP53BP1^ is changed. Nonetheless, this data suggests that the AUG^GFP^ initiation site is not the source of the protein band observed on SEP-GFP immunoblots.

**Figure 5:**
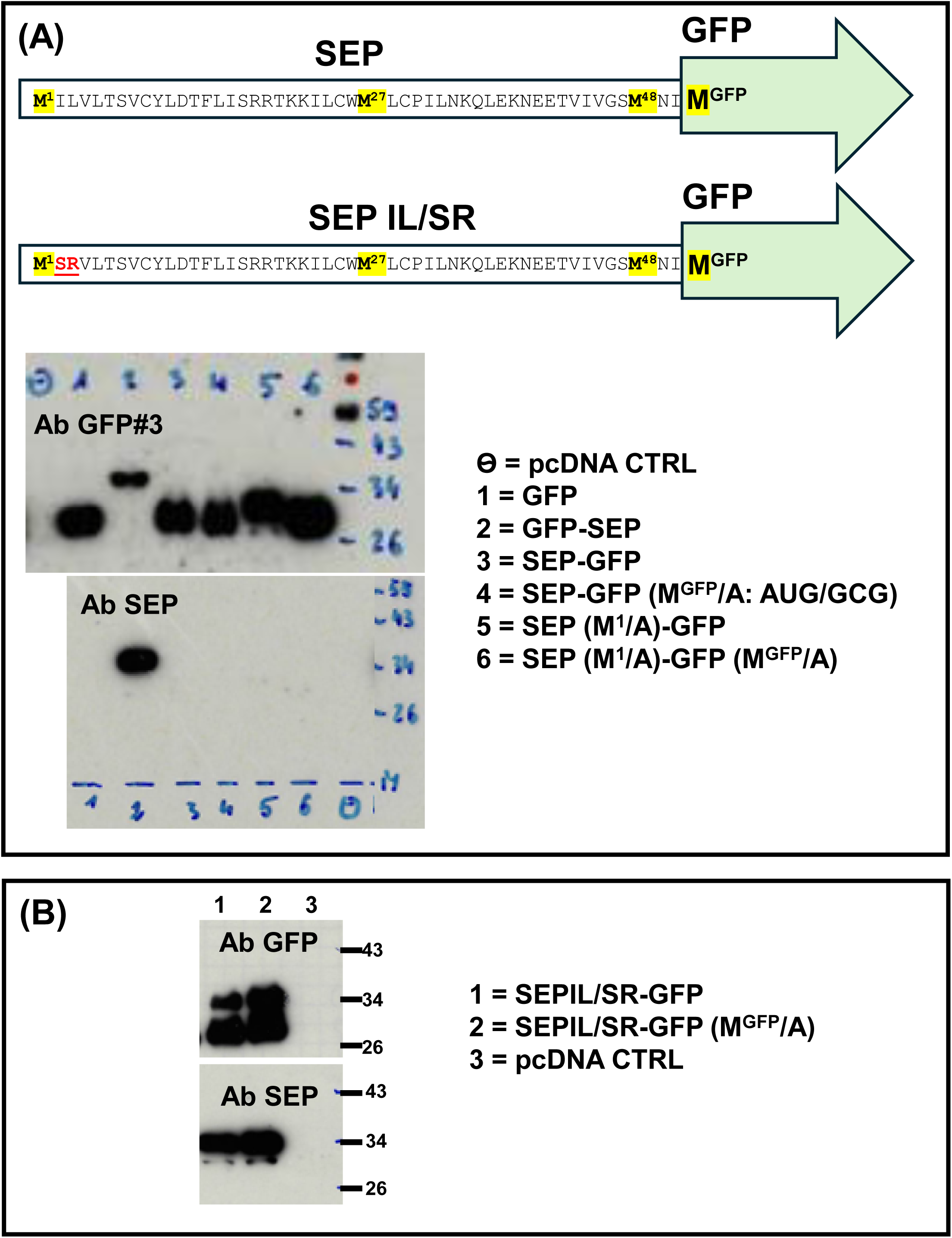
Expression profiles of SEP-GFP mutants within the SEP ORF. (A). The upper panel depicts the SEP-GFP fusion, the position of the IL/SR mutations and the alternative AUG start sites (M) within the SEP ORF (and hence fused to GFP). The lower panels are immunoblots performed with the polyclonal GFP#3 and SEP Abs using extracts prepared from HEK293T cells transfected with the constructs indicated on the right. (B). Immunoblots performed with the polyclonal GFP#3 and SEP Abs on extracts prepared from HEK293T cells transfected with the constructs indicated on the right.

The N-terminal of SEP^53BP1^ is largely non-polar (**MILVL**TS**V**C**YL**…) although it does not score as a signal sequence using the SignalP - 4.1, SignalP 6.0, TargetP-2.0 and Phobius analytical tools. Nonetheless, we have several observations suggesting that the protein interacts with membrane components in the cell, namely, co-localisation with the Golgi (Figure 4C), co-localisation with the endosome marker EEA1 (thesis Inchingolo, https://archive-ouverte.unige.ch/unige:164700) and its presence in extracellular exosomes (thesis Inchingolo, https://archive-ouverte.unige.ch/unige:164700). We therefore changed the I^2^/L^3^ positions to the polar S^2^ and charged R^3^ (SEPIL/SR). This SEP(IL/SR)-GFP construct produced two bands on anti-GFP#3 immunoblots (Figure 3C lane 4 and Figure 5B). The band co-migrating with GFP was present, but we also observed a new slower migrating band (migrating slightly faster than the GFP-SEP band: compare lanes 2 and 4 in the right-hand panel of Figure 3C) that was now also detected by the anti-SEP Ab (Figure 5B). This pattern of expression was not perturbed in the background M^GFP^/A (AUG/GCG mutation of the GFP start codon) suggesting that initiation events at the GFP start codon play little role in the observed expression profiles (Figure 5B).

With regards to intracellular localisation the behaviour of the SEP(IL/SR)-GFP construct was also intriguing. Whereas SEP-GFP gave a diffuse cellular staining reminiscent of GFP, and neither showed any co-localisation with the Golgi (Supplemental Figure S2), the staining observed in cells expressing the IL/SR mutant was both diffuse and punctate with the latter co-localising with the Golgi marker (Supplemental Figure S2). This might suggest that the two proteins observed when this construct is expressed accumulate in different intracellular compartments. The longer form, that presumably is the non-processed protein (see below), is co-localising with the Golgi whereas the shorter, and presumably cleaved form, behaves like GFP. This leads us to propose that SEP^53BP1^ acts as a Golgi target sequence and can function as such when placed N- or C-terminal.

The data suggest a post-translational processing event on the SEP-GFP protein that is coupled to the non-polar nature of the proteins N-terminal. However, to confirm this, we turned to proteolysis (trypsin was selected) coupled to mass spectrometry (mass spec) (Figure 6). The analysis (performed at the FGCZ, Zurich) focused on identifying tryptic peptides characteristic of the N-terminal region of SEP-GFP and SEP(IL/SR) and the C-terminal region of GFP-SEP (Figure 6A: boxed in red). Proteins were recovered from transfected cell extracts using the GFP#3 Ab and Protein G magnetic beads and the enrichment was confirmed on immunoblots using a fraction of the total beads (Figure 6B). In the initial study, the tryptic digestion was performed directly on the beads. Peptides covering all of GFP-SEP were detected, including the two C-terminal peptides within the SEP^53BP1^ region and the N-terminal GFP fragment covering aas 5-27 (boxed in red in Figure 6A). It should be noted that this analysis cannot confirm the exact N-terminal sequence since the four terminal amino acids MVSK would not be detected (although the presence of the tryptic peptide 5-27 is consistent with the presence of the K^4^ residue). In SEP-GFP, the N-terminal peptide covering aas 1-18 was not detected whereas in the IL/SR mutant the peptide covering aas 4-18 was observed. Because the analysis “on the bead” in the IL/SR background was looking at a mixed protein population we decided to refine the analysis by isolating the immunoselected protein bands from Coomassie stained gels (Figure 6C). We focused on the SEP-GFP band, and the two bands observed in the IL/SR mutant (indicated as U and L in Figure 6C). The analysis confirmed the presence of the N-terminal tryptic peptide 4-18 in the upper band (U) but a clear N-terminal truncation of 37 aas in both the mutant lower (L) band and the parent SEP-GFP protein (Figure 6C).

**Figure 6:**
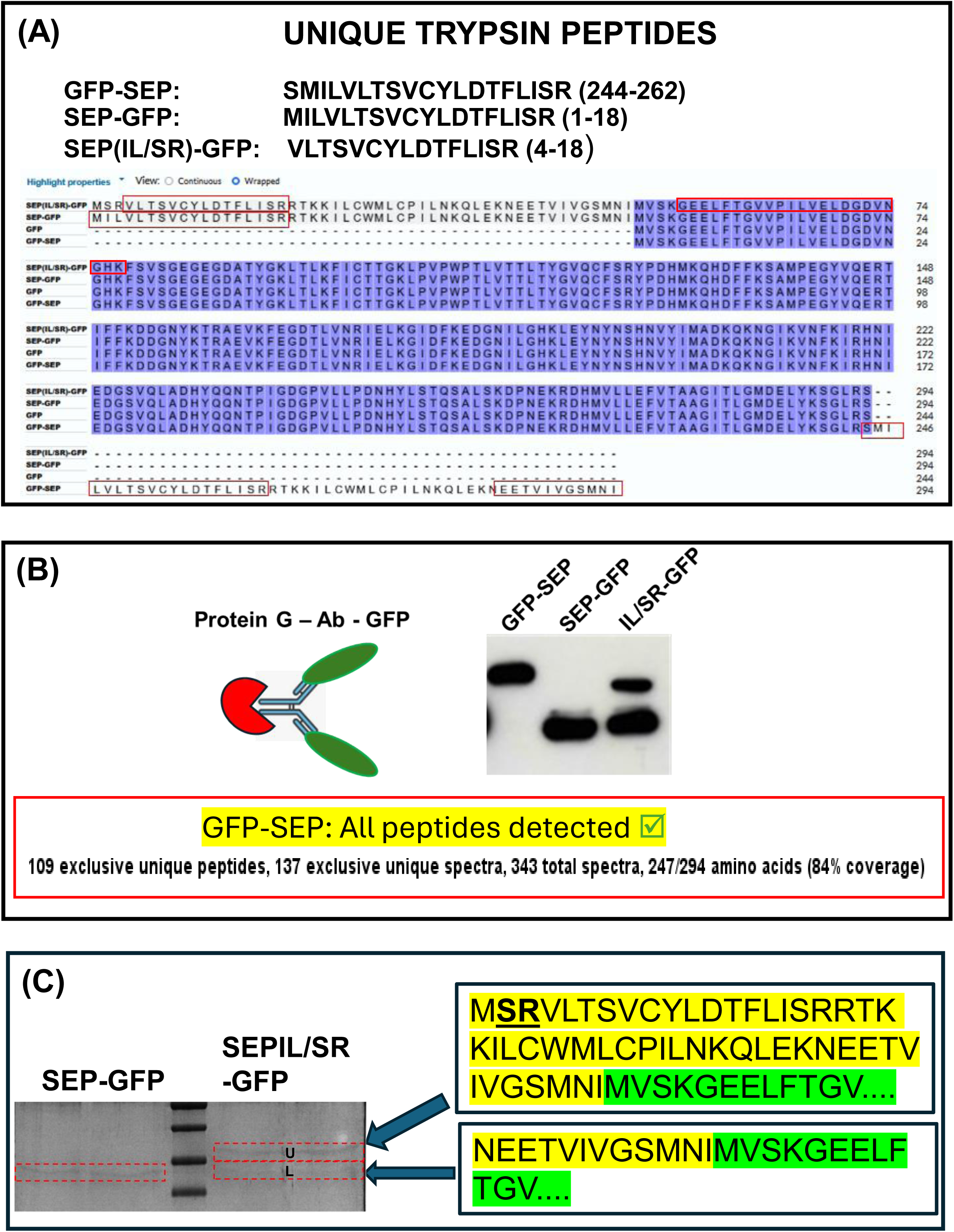
Mass-spec analysis of the tryptic fragments generated from the GFP-SEP, SEP-GFP and SEP(IL/SR)-GFP proteins. (A). An alignment of all proteins with tryptic peptides characteristic of the N-terminal of the SEP-GFP constructs and the GFP-SEP construct indicated. (B). The fusion proteins were bound to a Protein G magnetic bead via the GFP#3 Ab. Before sending for mass-spec analysis a small fraction of the beads was treated with sample buffer and analysed by immunoblotting with the same GFP#3 Ab (right-hand column). Proteins were trypsinised directly on the beads. All peptides for the GFP-SEP protein were detected. (C). Enrichment was performed as in (B) with the SEP-GFP and IL/SR mutant. Proteins were eluted from the beads with glycine buffer pH2 and loaded onto a preparative 12% PAGE gel. Bands were visualised by Coomassie staining. The gel slices corresponding to SEP-GFP and the upper (U) and lower (L) bands of the IL/SR were sent for trypsin mass-spec analysis. The determined N-terminal sequences are indicated on the right with the SEP^53BP1^ region highlighted in yellow and the beginning of GFP highlighted in green.

In conclusion, when placed N-terminal to a reporter the majority of the SEP^53BP1^ sequence is removed by a post-translational processing event. This event is coupled to the nature of the N-terminal sequence as it is limited, but not totally blocked, by mutations in this region. This cleavage does not occur when the SEP^53BP1^ sequence is placed C-terminal. Nonetheless, this latter construct, GFP-SEP, exhibits some unusual properties. It is not detected by several commercial monoclonal Abs and the GFP fluorescence appears to be quenched in transfected cells. Initially we suspected that N-terminal processing might also be occurring on GFP-SEP because small N-terminal truncations (∼8 aas) have been reported to impact negatively on intracellular fluorescence (40). However, the mass-spec data indicate that this is not the case (Figure 6). Since the protein associates with several intracellular membranes, including the Golgi and endosomes, we asked if some of the intracellular turnover was taking place in lysosomes. Employing inhibitors of the proteasome (MG132) and two inhibitors of lysosomal activity (chloroquine and bafilomycin) we followed GFP-SEP decay in cycloheximide (CHX) treated HEK293T cells (Supplemental Figure S3). In the presence of CHX the decay of GFP-SEP appears to be biphasic, with a rapid initial drop and a subsequent stabilisation. This may suggest two distinct intracellular pools of the protein. The results also reveal that only the proteasome inhibitor stabilises GFP-SEP, hence, little or no turn-over is occurring in lysosomes.

### Membrane association

As with the translational regulation of SEP expression, the behaviour of the GFP fusion proteins leaves us with a conundrum wrapped in an enigma. Previous immunofluorescent studies on fixed cells with the anti-SEP Ab suggested association with intracellular endosomes, although we have to admit that the antibody was far from ideal and gave high background signals (thesis Inchingolo, https://archive-ouverte.unige.ch/unige:164700). In a similar vein, SEP could be recovered in the medium and differential centrifugation studies placed at least a fraction of it in exosomes (thesis Inchingolo, https://archive-ouverte.unige.ch/unige:164700). The removal of a major part of the N-terminus on the SEP-GFP protein also remains curious. Clearly the hydrophobic amino acids 2 and 3 are important although their mutation to polar/charged amino acids did not completely block cleavage and this was not improved by the further mutation of the next two hydrophobic amino acids (producing an N-terminal sequence MSRDH….: Supplemental Figure S4). This suggests that another determinant resides within the body of the microprotein. We suspect that this may be the same signal that directs GFP-SEP to the Golgi. Cleavage is rapid and seems to involve a progressive trimming of the N-terminal SEP sequence (Figure 3B).

### The α4-SEP^53BP1^ interaction does not affect proteasome/immunoproteasome activity

Interactome studies indicated that SEP^53BP1^ interacted with multiple subunits of the proteasome. In particular, the interaction with the C-terminal tail of the α4 subunit of the 20S proteasome barrel scored high in our yeast-2-hybrid assay (23). We investigated if this interaction modulated proteasome activity. We employed lentiviral vectors to establish cell lines expressing SEP^53BP1^ as we were worried that transfection procedures themselves could impact on the readout and that in some of our cell lines, notable THP-1, transfection efficiency is poor. Starting with the HeLa-SEP^53BP1^ transduced cell line, in which SEP^53BP1^ expression was high, we reproduced our earlier published work (that used transient expression in transfected cells) showing co-sedimentation of SEP^53BP1^ and α4 (Figure 7A). Furthermore, both proteins showed a shift in the gradient in the presence of ATP consistent with the ATP-driven formation of the 26S proteasome subunit (41). However, the proteasome activity assayed in cell extracts was essentially indistinguishable in the transduced and HeLa parent cell lines (Figure 7B). We had observed low endogenous expression of SEP^53BP1^ in THP-1 cells (23). This is a human monocyte cell line in which immunoproteasome activity is high (42) (see below). Therefore, we next explored if SEP^53BP1^ expression could modulate the activity of the immunoproteasome. For this we used the same lentiviral vectors to establish THP-1 cells which we hoped would over-express SEP^53BP1^. The initial screening of the transduced cells revealed low SEP^53BP1^ transcription (the primer set was specific for the transduced SEP^53BP1^ cistron), and undetectable protein (the amount of cell extract loaded onto the gels, between 20-40 μg, was based upon the amounts used to observe the transiently expressed protein). In a second screening, we selected cells giving a high GFP signal (the transduced gene expresses an EMCV based bicistronic mRNA expressing GFP). We could now detect a SEP^53BP1^ transcript, albeit at levels lower than that observed in the transduced HeLa cells (Figure 8A). However, we were still unable to observe the protein in the transduced cells (again using 20-40 μg of protein extracts, quantities that do not permit detection of the endogenous protein), and it only became detectable when these cells were treated with MG132 (Figure 8B), a result consistent with a rapid proteasome-mediated degradation. Nonetheless, this was not an optimal experimental system in which to monitor the impact of SEP^53BP1^ on immunoproteasome activity.

**Figure 7:**
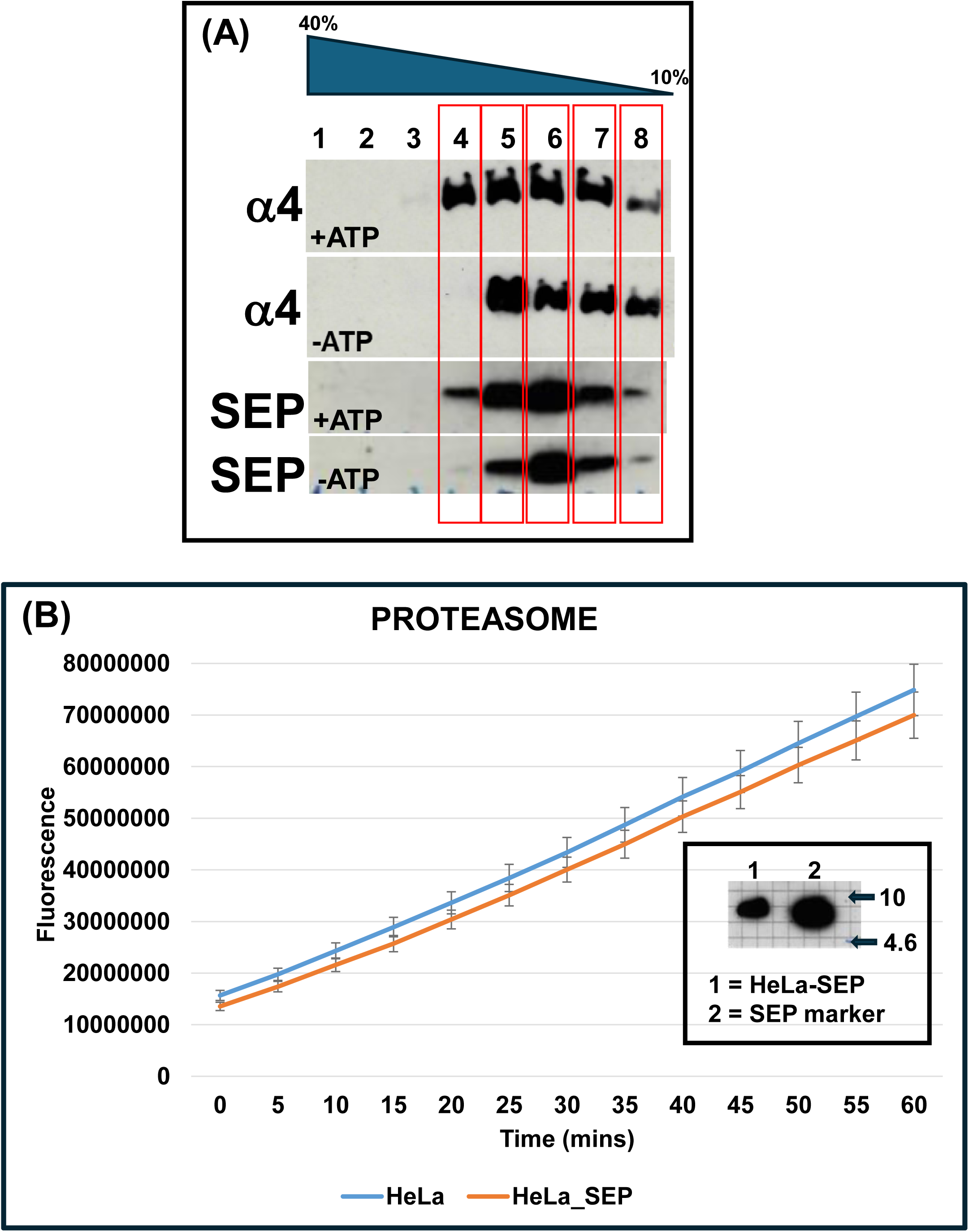
The impact of SEP^53BP1^ expression on proteasome activity. (A) Glycerol gradient analysis of lentiviral vector transduced HeLa cells stably expressing SEP^53BP1^. Cells were lysed in a hypotonic buffer with/without ATP. The ATP+ extracts were fractionated on gradients also containing ATP. Fractions were recovered as indicated and analysed by immunoblotting for the presence of α4 and SEP^53BP1^. (B) Proteasome assays were performed on hypotonic cell extracts prepared from transduced HeLa cells expressing SEP^53BP1^ and the parent HeLa cell line, using the Abcam Proteasome Activity Kit. Triplicate time points were analysed and plotted graphically as the mean and standard deviation (SD). The insert is an immunoblot of transduced HeLa cells expressing SEP^53BP1^ (SEP). A size marker is provided by a SEP^53BP1^ synthetic peptide.

**Figure 8:**
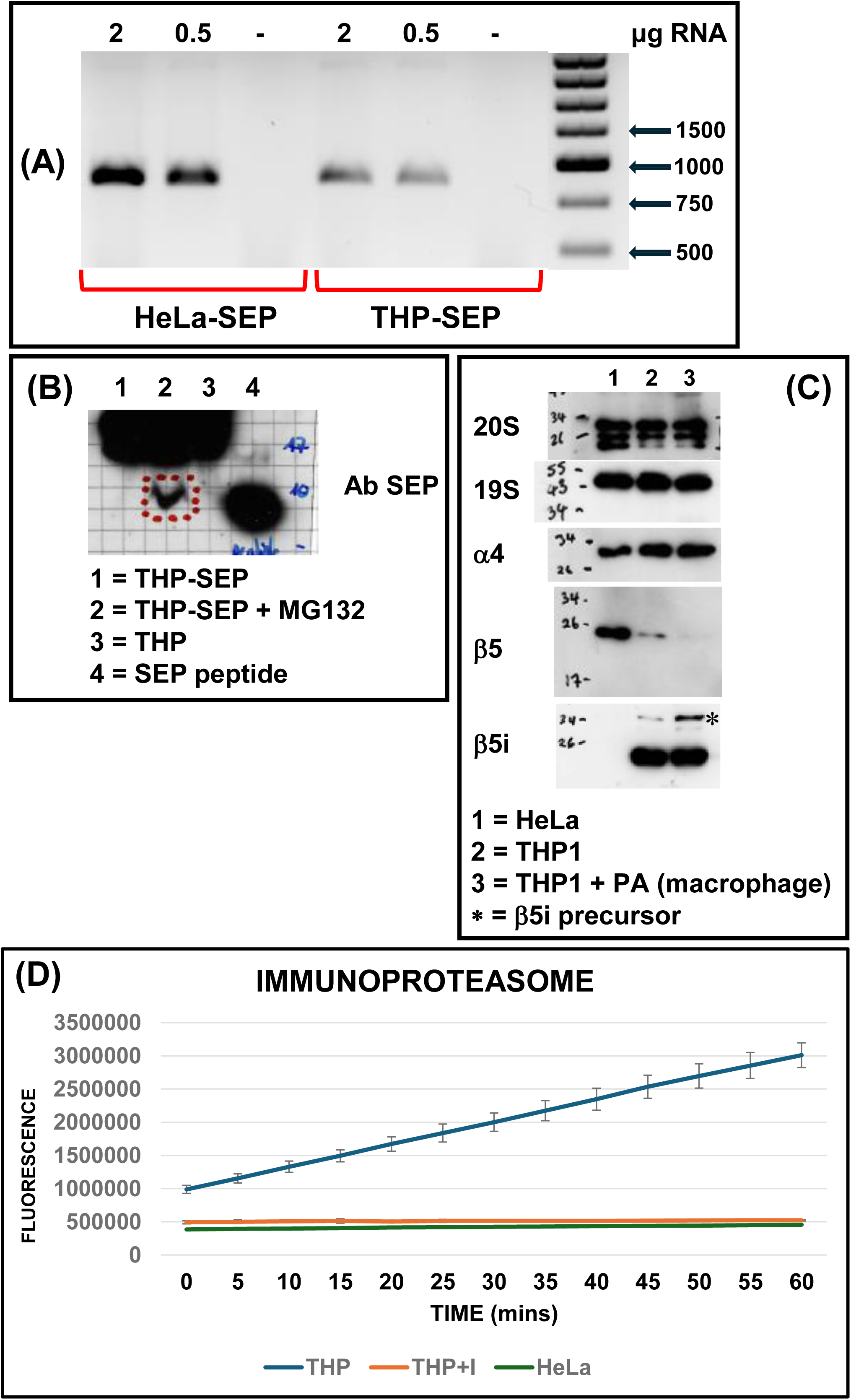
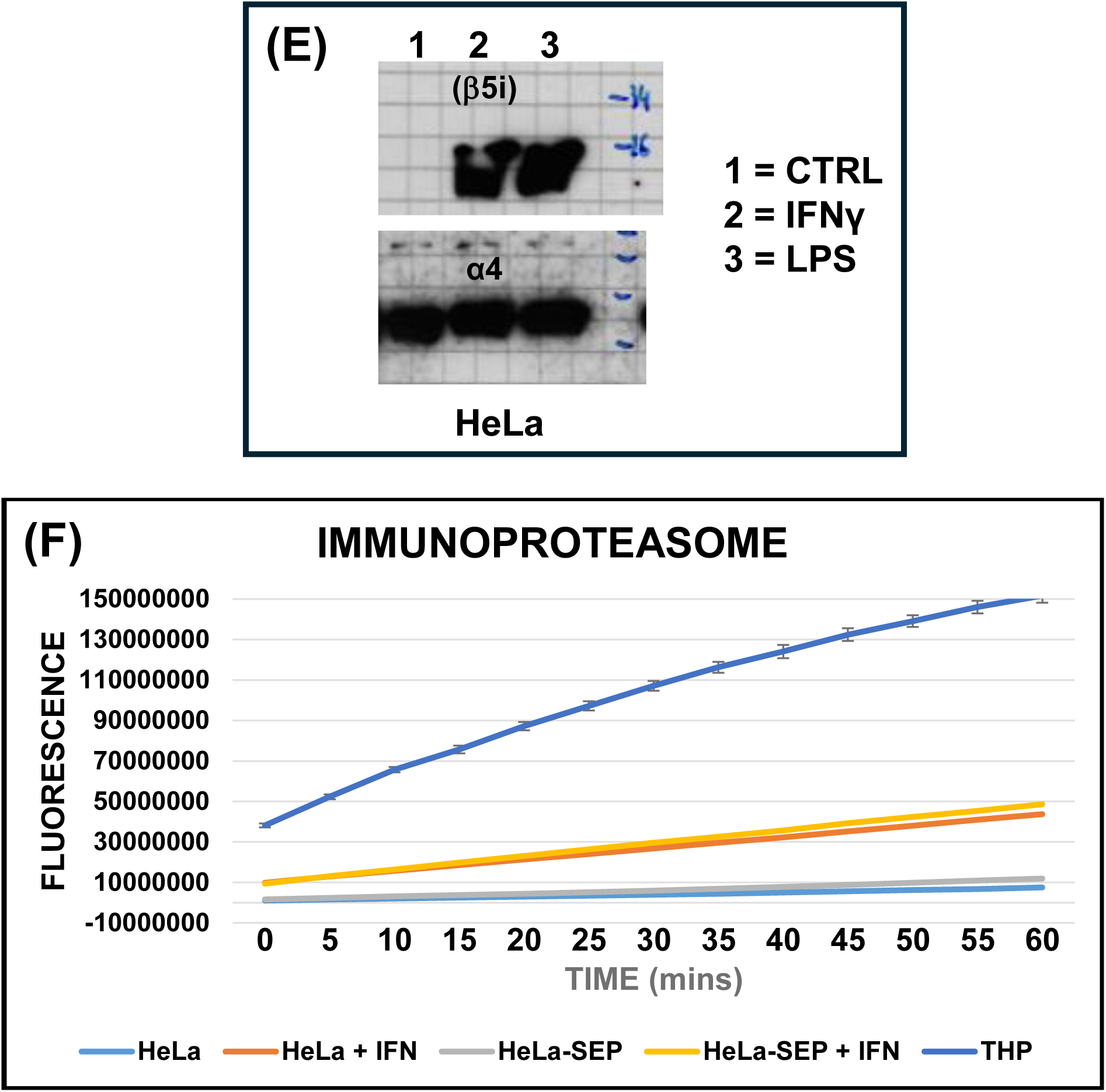
Immunoproteasome activity in HeLa and ThP-1 cells. (A). RT-PCR analysis of SEP^53BP1^ mRNA expression in the transduced cell lines. The starting material was total cell RNA (quantities are indicated above the panel). (B). A SEP^53BP1^ immunoblot prepared from THP-1 transduced cells. A size marker is provided by the SEP^53BP1^ synthetic peptide. (C). Immunoblots performed on HeLa, THP1 and macrophages (phorbol 12-myristate 13-acetate (PA) differentiated THP-1 cells) for specific proteasome (β5), immunoproteasome (β5i) and shared markers (20S. 19S. α4). (D). Assays were performed on cell extracts from HeLa and THP-1 cells using the immunoproteasome substrate Ac-Ala-Asn-Trp-AMC (AdipoGen) plus or minus the specific immunoproteasome inhibitor ONX 0914 (AdipoGen). Each time point was monitored in triplicate and plotted graphically as the mean plus the SD. (E). An immunoblot monitoring immunoproteasome induction in HeLa cells lines treated with interferon gamma (IFNγ) or bacterial lipopolysaccharide (LPS) as determined by the expression of β5i (also called LMP7). (F). Immunoproteasome assays performed on the indicated cell lines. (C). This is the same data as presented in panel (B) excluding the THP-1 cell line.

Unlike THP-1 cells, HeLa cells do not express the characteristic subunits of the immunoproteasome 20S barrel, namely β1i, β2i and β5i (Figure 8C) and exhibit activity comparable to that observed in THP-1 cells in the presence of a specific immunoproteasome inhibitor (Figure 8D). However, expression of the βi subunits can be triggered by treatment with cytokines such as interferon gamma (IFNγ) (43) or endotoxins such as bacterial lipopolysaccharides (LPS) (Figure 8E) (44). The IFNγ induced expression of β5i correlated with a significant increase in immunoproteasome activity in the HeLa cells, albeit at levels lower than that observed in THP-1 cells (Figure 8F). However, the immunoproteasome activity profile was unperturbed in the transduced HeLa cell lines expressing SEP^53BP1^ (Figure 8F).

Thus, although the SEP^53BP1^ interacts with components of the proteasome/immunoproteasome we have been unable to demonstrate that its presence alters the proteolytic activity of either. We have not explored the possibility that it may be altering the nature of the polypeptides generated and subsequently presented on the cell surface by the MHC complex, the immunopeptidome (45).

## CONCLUSIONS

We have accumulated a considerably amount of data on the properties of the SEP^53BP1^ microprotein, its unusual translational regulation, its intracellular and possibly extracellular location. However, its function remains elusive. It is possible that this will only manifest itself in the context of the whole animal. However, this is not the first report of a microprotein arising from an altORF carrying a Golgi targeting sequence. A 37 aas altORF within the mRNA of the centromere protein CENP-R was recently reported to also serve as a Golgi targeting microprotein. Like SEP^53BP1^, it could direct GFP fusions to the cytoplasmic surface of the Golgi (46). It is conserved in primates (less so in other mammals), and the authors identified a consensus sequence containing a critical cysteine residue. However, this consensus sequence is not present in SEP^53BP1^, suggesting a different mode of targeting. The Golgi apparatus represents a central relay site for intracellular, and ultimately extracellular, membrane trafficking (47). This close association between SEP^53BP1^ and the Golgi may also explain our observed association with both endosomes and exosomes. As our lab has now closed, it is hoped that the information provided in this manuscript will form the foundation of future studies by parties interested in microprotein expression/function.

## MATERIALS AND METHODS

### Cell culture and transfection

HEK293T, HeLa, MCF7, MCF10A, THP-1 and Raji cells were grown at 37°C in a humidified 5% CO_2_ chamber. Cells were cultured in Dulbecco’s modified Eagle’s medium (DMEM) supplemented with 1% penicillin/streptomycin (P/S) (Gibco) and 10% fœtal bovine serum (FBS, Brunschwig) for HEK293T, HEK293, MDA-MB-231, and HeLa cells. MCF7 (ATCC, HTB-22) were cultured DMEM F-12 medium (Gibco), 1% P/S and 10% FBS supplemented with 10 µg/mL human recombinant insulin and 0.5 nM estradiol. MCF10A (ATCC, CRL-10317) were cultured DMEM F-12 medium containing 1% P/S, 5% horse serum heat inactivated (HS, Brunschwig), 10µg/mL Epidermal Growth Factor (EGF), 1 µM Dexamethasone, and 5µg/ml human recombinant insulin. THP-1 and Raji cells (kind gift of Prof. Walter Reith, UNIGE) were cultured in RPM1 1640 1X (+L-Glutamine) supplemented with 0.05mM 2-mercaptoethanol and 10% fœtal bovine serum. MRC-5 cells were grown in Eagle’s Minimum Essential Medium (MEM; Sigma) supplemented with 10% FBS, 1% P/S, 1X amino acids and 1mM sodium pyruvate. HCT-116 cells were cultured in McCoy’s 5a Medium Modified (Gibco) supplemented with 10% FBS and 1% P/S.

Transfections of HEK293T cells were performed using Lipofectamine 3000 (Invitrogen) when the cells were 70-80% confluent. Eight hours post-transfection, the medium was replaced with normal growth medium, and lysates were usually prepared at 24h post-transfection.

### Drug Treatment

Transfected HEK293T cells were treated with MG132 (10 μM), chloroquine (100 μM) or bafilomycin (100 nM) at 16 hrs post-transfection. DMSO treated cells served as a control. For protein half-life assays cells were lysed in CSH buffer.

### DNA cloning

Unless stated otherwise, all clones were prepared in a pcDNA3 backbone. Mutations, fusions and deletions were introduced by PCR. The oligos employed are listed in the Supplementary Table 1. The EMCV, HCV and CrPV IRES plasmid backbones have been described previously (48). The mCherry Golgi marker construct (pmCherry-N1-GalT) was obtained from Addgene. The GFP coding region was subcloned using the plasmid pT8MycGFP-HX (a kind gift of D. Soldati-Favre, University of Geneva, Switzerland) as a template (49). All fusions to SEP^53BP1^ retained the original AUG start codons on each ORF. Mutations and deletions were introduced by PCR (see oligo list).

### RT-PCR

RNA was prepared directly from transfected cell pellets using the TRIzol reagent (Invitrogen). It was treated with DNase (DNA free; Ambion), and quality was checked on an Agilent 2100 bioanalyzer. Semi-quantitative RT-PCR was performed starting with 100 ng of total cell RNA using the QIAGEN OneStep RT-PCR Kit (in the RT minus control samples were heated to 95°C for 5 mins prior to starting. The number of amplifications cycles was first optimised for the primer set and corresponded to the exponential phase. For the GFP and GFP fusion proteins the primer set corresponded to the SP6/T7 sequences present on the pcDNA3 transfected plasmids.

### Western blot

Cytoplasmic extracts were prepared in CSH buffer (50 mM Tris-Cl pH 7.5, 250 mM NaCl, 1 mM EDTA, 0.1% (v/v) Triton X-100). Alternatively, whole cell extracts were prepared by resuspending the cell pellet in either X2 sample buffer (125 mM Tris-HCl pH 6.8, 20% glycerol, 10% (v/v) β-mercaptoethanol, 5% SDS, 0.025% (w/v) bromophenol blue) for SDS-PAGE, or X2 sample buffer Novex (450mM Tris H-Cl pH 8.45, 12% Glycerol, 4% SDS, 0.00075% Coomassie Blue G, 10% β-mercaptoethanol) for Tricine gels (Invitrogen).

Protein concentrations were determined by Bradford (Cytoskeleton, USA). Twenty μg of protein was resolved on the polyacrylamide gels and electro-transferred to PVDF membranes (semi-dry transfer foe SDS-PAGE gels and wet transfer for the Tricine gels).

Antibodies used in this study were: anti-SEP^53BP1^ (23), anti-GFP#1 (Roche mouse monoclonal #C755C12), anti-GFP#2 (Abcam mouse monoclonal #ab1218), anti-GFP#3 (Proteintech rabbit polyclonal antibody, #50430-2-AP), anti-HA (Covance clone 16B12), anti-FLAG (M2 antibody, Sigma), anti-actin (Millipore, #MAB1501), anti-β5 (Santa Cruz Biotechnologies, sc-393931), anti-β5i (ENZO, BML-PW8845), anti-α4 (PSMA7, MCP72, Santa Cruz Biotechnologies, sc-58417), anti-20S/anti-19S proteasome (Enzo) and goat anti-mouse or rabbit HRP secondary antibodies (Bio-Rad). The Anti-MYC tag was a gift from Prof. Dominique Soldati (University of Geneva, Switzerland). Immunoblots were developed using the WesternBright™ Quantum (Advansta) and quantitated using Image Lab (Biorad).

### Glycerol gradients

Glycerol gradients were prepared as previously described(50). HEK293T cells were lysed in hypotonic lysis buffer (see below) containing 2mM ATP.

Cell extracts were loaded onto a 10–40% linear glycerol gradient in 100 mM KCl, 5 mM MgCl_2_, and 20 mM HEPES pH 7.4 prepared in an SW60 tube. Gradients were centrifuged for 16 hr at 30,000 rpm in a SW60 rotor at room temperature. After centrifugation, 10 X 400 μL fractions were collected from the bottom of the tube. Proteins were recovered by methanol/chloroform precipitation, resuspended directly in 40μL of X2 sample buffer and analysed by western blotting.

### Live cell imaging

GFP and GFP-fusion constructs were transiently expressed in either HEK293T or HeLa cells. Live cell confocal images were collected at 18 hrs post-transfection on a STELLARIS 5 confocal scanning microscope equipped with a HC PL APO CS2 63x/1.40 oil objective. Pictures were analysed using the ImageJ and Imaris software.

### Viral particles production and transduction

HEK293T cells at 80% confluence were transfected with the second-generation packaging and envelope vectors (pWPI: https://www.addgene.org/12254/). Viral particles were filtered through a 0.45 μm filter at 48 h post-transfection, and used to transduce HeLa and THP1 cells. Five days later, GFP expressing cells were sorted by flow cytometry.

### Proteasomal/Immunoproteasome activity assays

The proteasomal activity assay was performed on either transfected cells expressing SEP^53BP1^ (with an empty vector control) or lentivirus transduced cells. Proteasome activity was measured using the Proteasome Activity kit (ab107921, Abcam) following the manufacturer’s protocol. MG-132 (Sigma) treatment served as a control to differentiate between 26S proteasome activity and other protease activities present in the extract. The reaction was followed for 90 minutes, and fluorescence was measured on a fluorometric microplate reader (SpectraMax Paradigm) equipped with an Ex/Em: 345/445nm filter. Timepoints were performed in triplicate. The immunoproteasome assay was performed in identical buffer conditions but with the fluorogenic peptidyl substrate Ac-Ala-Asn-Trp-AMC (final concentration 10 μM) (AdipoGen) and the specific inhibitor ONX 0914 (10 μM) (AdipoGen) (42,43,51).

### Protein Characterisation

Proteins were transiently expressed in HEK293T cells. Extracts were prepared in RIPA buffer (150 mM NaCl, 1% deoxycholate, 1% Triton X-100, 0.1% SDS, 10 mM Tris pH 7.8) and proteins recovered by immunoprecipitation using the anti-SEP^53BP1^ Ab prebound to Protein-A magnetic beads (ThermoFisher).

Immunoselected proteins resolved on TRICINE gels were subsequently stained with ReadyBlue™ Protein Gel Stain (Merck). Protein bands were excised and sent for tandem mass spectra analyse at the Functional Genomics Center Zurich. All MS/MS samples were analysed using PEAKS Studio (Bioinformatics Solutions, Waterloo, ON Canada; version 10.5 (2020-02-19) assuming the digestion enzyme trypsin and compared directly with the known primary sequence of the transfected proteins. PEAKS Studio used a fragment ion mass tolerance of 0,020 Da and a parent ion tolerance of 10,0 PPM. Oxidation of methionine and carbamidomethyl of cysteine were specified in PEAKS Studio as variable modifications. Scaffold (version Scaffold_5.3.3, Proteome Software Inc., Portland, OR) was used to validate MS/MS based peptide and protein identifications. Peptide identifications were accepted if they could be established at greater than 59,0% probability to achieve an FDR less than 0,1% by the Percolator posterior error probability calculation (52). Protein identifications were accepted if they could be established at greater than 5,0% probability to achieve an FDR less than 1,0% and contained at least 2 identified peptides. Protein probabilities were assigned by the Protein Prophet algorithm (53).

## Supporting information

Supplemental Results

## Oligonucleotide list

SV40_IntronHIII(+)

(5’-3’): TCT AAG CTT CAA GCT CCT CGA GGA ACT GAA AAA CCA GAA AG

SV40_IntronHIII(-)

(5’-3’): TCT AAG CTT CAG CTT TTA GAG CAG AAG TAA CAC TTC CGT AC

GFP_D50aa

(5’-3’): TCT AAG CTT ACC ATG GGC AAG CTG CCC GTG CCC TGG CCC ACC CTC G

SEP_Nterm_MUT4aas

(5’-3’): TCT AAG CTT ACC ATG AGT CGG ATC ACA CTT CAG TAT GCT ATC TCG ACA CCT TC

FLAG_MYC_STAB_FS(-)

(5’-3’): GCC GTT ACT AGT GGA TCC TCT GGA TTT GTT TTT TTT GCG AGT C

EMCV_IRES_Nco(-)

(5’-3’): CTT CCA TGG TAT TAT CGT GTT TTT CAA AGG AAA ACC AC

HCV_IRES_HIII(+)

(5’-3’): CTC AAG CTT GCC ATA GCG TTA GTA TGA GTG TCG TAC AGC

HCV_IRES_Nco(-)

(5’-3’): AGC CCA TGG TGC ACG GTC TAC GAG ACC TCC CGG GGC

CrPV_IRES_HIII(+)

(5’-3’): CTC AAG CTT GAA TTC AAA GCA AAA ATG TGA TCT TGC TTG TAA ATA C

SEP_Nco(+)

(5’-3’): CTC CCA TGG TTC TGG TTC TCA CTT CAG TAT GCT ATC TCG AC

SEP.AUG_402.GCGHIII(+)

(5’-3’): CCC AAG CTT AGG ATG ATT CTG GTT CTC ACT TCA GTG CGC TAT CTC GAC AC

SEP.GCG_402.GCGHIII(+)

(5’-3’): CCC AAG CTT AGG GCG ATT CTG GTT CTC ACT TCA GTG CGC TAT CTC GAC AC

V3_FLAG.MYC.STAB(+)

(5’-3’): GCC TGA AAG CCA GGT TCT AGA GGA TGG TTG GAG CAG TTG CCG ACT ACA AAG ACG

V3(-)

(5’-3’): CAT CCT CTA GAA CCT GGC TTT CAG GC

SEP-GFP.GCG_AUG3LEU(+)

(5’-3’): GAC GGT AAT AGT GGG TTC ACT AAA CAT TGC GGT GAG CAA GGG CGA GGA GC

SEP-GFP.GCG_AUG3LEU(-)

(5’-3’): GCT CCT CGC CCT TGC TCA CCG CAA TGT TTA GTG AAC CCA CTA TTA CCG TC

SEP-GFP_AUG3LEU(+)

(5’-3’): GAC GGT AAT AGT GGG TTC ACT AAA CAT TAT GGT GAG CAA GGG CGA GGA

SEP-GFP_AUG3LEU(-)

(5’-3’): GCT CCT CGC CCT TGC TCA CCA TAA TGT TTA GTG AAC CCA CTA TTA CCG TC

GCG_AUG2_SEP(+)

(5’-3’): GCA CAA AGA AAA TCC TGT GTT GGG CGT TGT GTC CAA TCC TGA ACA AAC

GCG_AUG2_SEP(-)

(5’-3’): GTT TGT TCA GGA TTG GAC ACA ACG CCC AAC ACA GGA TTT TCT TTG TGC

SEPwt_GFP_HIII (+)

(5’-3’): TCT AAG CTT ACC ATG ATT CTG GTT CTC ACT TCA GTA TGC TAT CTC GAC AC

SEP_IS_LR _GFP_HIII (+)

(5’-3’): TCT AAG CTT ACC ATG AGT CGG GTT CTC ACT TCA GTA TGC TAT CTC GAC ACC TTC

FLAG-MYC-STAB (+) (5’-3’): ATG GTT GGA GCA GTT GCC GAC TAC

FLAG-MYC-STAB (-) (5’-3’): TTT GCG AGT CCT TGC CGT CGC TC

SEPAUG_GCG HIII(+)

(5’-3’): TCT AAG CTT GCG ATT CTG GTT CTC ACT TCA GTA TGC TAT CTC GAC

SEP(+)

(5’-3’): ATG ATT CTG GTT CTC ACT TCA GTA TGC

GFP(-)

(5’-3’): CTT GGT CAG GCC GTG CTT GGA CTG

V3SEP_AUG-GCG(+)

(5’-3’): GCT GTA CAA GTC CGG ACT CAG ATC TGC GAT TCT GGT TCT CAC TTC AGT ATG C

V3SEP_AUG-GCG(-)

(5’-3’): GCA TAC TGA AGT GAG AAC CAG AAT CGC AGA TCT GAG TCC GGA CTT GTA CAG C

SEP_GFP_GCG(+)

(5’-3’): GAC GGT AAT AGT GGG TTC AAT GAA CAT TGC GGT GAG CAA GGG CGA GGA GCT GTT C

SEP_GFP_GCG (-)

(5’-3’): GAA CAG CTC CTC GCC CTT GCT CAC CGC AAT GTT CAT TGA ACC CAC TAT TAC CGT C

V3_FUS_FlagMyc+

(5’-3’): GGT TCT AGA GGA TGG TTG GAG CAG TTG CCG AC

GFP_Xba (-)

(5’-3’): CCC TCT AGA CTA AGA TCT GAG TCC GGA CTT GTA CAG C

50aa_FLAG_HA(-)

(5’-3’): CTG TCT AGA TTA TGG GTC GTC TCC GCC GCC TCC CGC

HindIII-FLAG-HA-50aa +ve

(5’-3’): CTC AAG CTT GAT ATG GTT GGA GCC GCA GTT GCC G

XbaI-FLAG-HA-50aa -ve

(5’-3’): GAG TCT AGA CTA TCC TGG GTC GTC TCC GCC GC

## Acknowledgements

We would like to thank Prof. Dominique Belin for helpful discussions.

## Author contributions

All authors performed the experiments. J.A.C designed the project, interpreted the data and wrote the article. J.A.C. supervised the project.

## References

1. Frith, M.C., Forrest, A.R., Nourbakhsh, E., Pang, K.C., Kai, C., Kawai, J., Carninci, P., Hayashizaki, Y., Bailey, T.L. and Grimmond, S.M. (2006) The abundance of short proteins in the mammalian proteome. PLoS Genet, 2, e52.

2. Cardon, T., Fournier, I. and Salzet, M. (2021) Shedding Light on the Ghost Proteome. Trends Biochem Sci, 46, 239–250.

3. Ingolia, Nicholas T., Lareau, Liana F. and Weissman, Jonathan S. (2011) Ribosome Profiling of Mouse Embryonic Stem Cells Reveals the Complexity and Dynamics of Mammalian Proteomes. Cell, 147, 789–802.

4. Tong, G. and Martinez, T.F. Ribosome profiling reveals hidden world of small proteins. Trends in Genetics.

5. Yang, H., Li, Q., Stroup, E.K., Wang, S. and Ji, Z. (2024) Widespread stable noncanonical peptides identified by integrated analyses of ribosome profiling and ORF features. Nature Communications, 15, 1932.

6. Azam, S., Yang, F. and Wu, X. (2025) Finding functional microproteins. Trends in Genetics, 41, 107–118.

7. Saghatelian, A. and Couso, J.P. (2015) Discovery and characterization of smORF-encoded bioactive polypeptides. Nature chemical biology, 11, 909–916.

8. Mumtaz, M.Ali, S. and Couso, Juan P. (2015) Ribosomal profiling adds new coding sequences to the proteome. Biochemical Society Transactions, 43, 1271–1276.

9. Guo, Z.-W., Meng, Y., Zhai, X.-M., Xie, C., Zhao, N., Li, M., Zhou, C.-L., Li, K., Liu, T.-C., Yang, X.-X., et al. (2019) Translated Long Non-Coding Ribonucleic Acid ZFAS1 Promotes Cancer Cell Migration by Elevating Reactive Oxygen Species Production in Hepatocellular Carcinoma. 10.

10. Whited, A.M., Jungreis, I., Allen, J., Cleveland, C.L., Mudge, J.M., Kellis, M., Rinn, J.L. and Hough, L.E. (2024) Biophysical characterization of high-confidence, small human proteins. Biophys Rep (N Y), 4, 100167.

11. Curran, J.A. and Weiss, B. (2016) What Is the Impact of mRNA 5’ TL Heterogeneity on Translational Start Site Selection and the Mammalian Cellular Phenotype? Front Genet, 7, 156.

12. Araud, T., Genolet, R., Jaquier-Gubler, P. and Curran, J. (2007) Alternatively spliced isoforms of the human elk-1 mRNA within the 5’ UTR: implications for ELK-1 expression. Nucleic Acids Research, 35, 4649–4663.

13. Carninci, P., Sandelin, A., Lenhard, B., Katayama, S., Shimokawa, K., Ponjavic, J., Semple, C.A., Taylor, M.S., Engstrom, P.G., Frith, M.C. et al. (2006) Genome-wide analysis of mammalian promoter architecture and evolution. Nat Genet, 38, 626–635.

14. Sandelin, A., Carninci, P., Lenhard, B., Ponjavic, J., Hayashizaki, Y. and Hume, D.A. (2007) Mammalian RNA polymerase II core promoters: insights from genome-wide studies. Nature Reviews Genetics, 8, 424.

15. Danino, Y.M., Even, D., Ideses, D. and Juven-Gershon, T. (2015) The core promoter: At the heart of gene expression. Biochimica et Biophysica Acta (BBA) - Gene Regulatory Mechanisms, 1849, 1116–1131.

16. Mauro, V.P. and Matsuda, D. (2016) Translation regulation by ribosomes: Increased complexity and expanded scope. RNA Biol, 13, 748–755.

17. Zhan, Y., Hu, Z., Lu, Z. and Lin, Z. (2025) The patterns of alternative TSS usage explain the highly heterogeneous landscape of 5’UTR lengths in eukaryotes. NAR Genom Bioinform, 7, lqaf152.

18. Dieudonne, F.X., O’Connor, P.B., Gubler-Jaquier, P., Yasrebi, H., Conne, B., Nikolaev, S., Antonarakis, S., Baranov, P.V. and Curran, J. (2015) The effect of heterogeneous Transcription Start Sites (TSS) on the translatome: implications for the mammalian cellular phenotype. BMC Genomics, 16, 986.

19. Mauro, V.P. and Edelman, G.M. (2002) The ribosome filter hypothesis. Proc Natl Acad Sci U S A, 99, 12031–12036.

20. Mauro, V.P. and Edelman, G.M. (2007) The ribosome filter redux. Cell Cycle, 6, 2246–2251.

21. Mirman, Z. and de Lange, T. (2020) 53BP1: a DSB escort. Genes & Development, 34, 7–23.

22. Dever, T.E., Ivanov, I.P. and Hinnebusch, A.G. (2023) Translational regulation by uORFs and start codon selection stringency.

23. Inchingolo, M.A., Diman, A., Adamczewski, M., Humphreys, T., Jaquier-Gubler, P. and Curran, J.A. (2023) TP53BP1, a dual-coding gene, uses promoter switching and translational reinitiation to express a smORF protein. iScience, 26, 106757.

24. Shi, X., Jiang, W., Yang, X., Li, Y., Zhong, X., Niu, J. and Shi, Y. (2024) TIR8 protects against nonalcoholic steatohepatitis by antagonizing lipotoxicity-induced PPARα downregulation and reducing the sensitivity of hepatocytes to LPS. Transl Res, 272, 68–80.

25. Zhang, Y., Lan, M., Liu, C., Wang, T., Liu, C., Wu, S. and Meng, Q. (2023) Islr regulates insulin sensitivity by interacting with Psma4 to control insulin receptor alpha levels in obese mice. Int J Biochem Cell Biol, 159, 106420.

26. Takebe, Y., Seiki, M., Fujisawa, J., Hoy, P., Yokota, K., Arai, K., Yoshida, M. and Arai, N. (1988) SR alpha promoter: an efficient and versatile mammalian cDNA expression system composed of the simian virus 40 early promoter and the R-U5 segment of human T-cell leukemia virus type 1 long terminal repeat. Mol.Cell Biol., 8, 466–472.

27. Nott, A., Meislin, S.H. and Moore, M.J. (2003) A quantitative analysis of intron effects on mammalian gene expression. RNA, 9, 607–617.

28. Nott, A., Le Hir, H. and Moore, M.J. (2004) Splicing enhances translation in mammalian cells: an additional function of the exon junction complex. Genes Dev, 18, 210–222.

29. Giorgi, C., Yeo, G.W., Stone, M.E., Katz, D.B., Burge, C., Turrigiano, G. and Moore, M.J. (2007) The EJC factor eIF4AIII modulates synaptic strength and neuronal protein expression. Cell, 130, 179–191.

30. Kozak, M. (1986) Point mutations define a sequence flanking the AUG initiator codon that modulates translation by eukaryotic ribosomes. Cell, 44, 283–292.

31. Rahim, G., Araud, T., Jaquier-Gubler, P. and Curran, J. (2012) Alternative splicing within the elk-1 5’ untranslated region serves to modulate initiation events downstream of the highly conserved upstream open reading frame 2. Mol Cell Biol, 32, 1745–1756.

32. Rethi-Nagy, Z., Abraham, E., Udvardy, K., Klement, E., Darula, Z., Pal, M., Katona, R.L., Tubak, V., Pali, T., Kota, Z. et al. (2022) STABILON, a Novel Sequence Motif That Enhances the Expression and Accumulation of Intracellular and Secreted Proteins. International journal of molecular sciences, 23.

33. Chu, V., Feng, Q., Lim, Y. and Shao, S. (2021) Selective destabilization of polypeptides synthesized from NMD-targeted transcripts. Mol Biol Cell, 32, ar38.

34. Monaghan, L., Longman, D. and Cáceres, J.F. (2023) Translation-coupled mRNA quality control mechanisms. 42, e114378.

35. Carrard, J. and Lejeune, F. (2023) Nonsense-mediated mRNA decay, a simplified view of a complex mechanism. BMB Rep, 56, 625–632.

36. Hertz, M.I. and Thompson, S.R. (2011) Mechanism of translation initiation by Dicistroviridae IGR IRESs. Virology, 411, 355–361.

37. Holbrook, J., Neu-Yilik, G., Gehring, N., Kulozik, A. and Hentze, M. (2006) Internal ribosome entry sequence-mediated translation initiation triggers nonsense-mediated decay. EMBO reports, 7, 722–726.

38. Isken, O., Kim, Y.K., Hosoda, N., Mayeur, G.L., Hershey, J.W. and Maquat, L.E. (2008) Upf1 phosphorylation triggers translational repression during nonsense-mediated mRNA decay. Cell, 133, 314–327.

39. Hanson, D.A. and Ziegler, S.F. (2004) Fusion of green fluorescent protein to the C-terminus of granulysin alters its intracellular localization in comparison to the native molecule. J.Negat.Results Biomed., 3, 2.

40. Li, X., Zhang, G., Ngo, N., Zhao, X., Kain, S.R. and Huang, C.-C. (1997) Deletions of the Aequorea victoria Green Fluorescent Protein Define the Minimal Domain Required for Fluorescence*. Journal of Biological Chemistry, 272, 28545–28549.

41. Orino, E., Tanaka, K., Tamura, T., Sone, S., Ogura, T. and Ichihara, A. (1991) ATP-dependent reversible association of proteasomes with multiple protein components to form 26S complexes that degrade ubiquitinated proteins in human HL-60 cells. FEBS Lett, 284, 206–210.

42. Vagapova, E., Burov, A., Spasskaya, D., Lebedev, T., Astakhova, T., Spirin, P., Prassolov, V., Karpov, V. and Morozov, A. (2021) Immunoproteasome Activity and Content Determine Hematopoietic Cell Sensitivity to ONX-0914 and to the Infection of Cells with Lentiviruses. Cells, 10, 1185.

43. Kim, S., Park, S.H., Choi, W.H. and Lee, M.J. (2022) Evaluation of Immunoproteasome-Specific Proteolytic Activity Using Fluorogenic Peptide Substrates. Immune network, 22, e28.

44. Reis, J., Guan, X.Q., Kisselev, A.F., Papasian, C.J., Qureshi, A.A., Morrison, D.C., Van Way, C.W., 3rd, Vogel, S.N. and Qureshi, N. (2011) LPS-induced formation of immunoproteasomes: TNF-α and nitric oxide production are regulated by altered composition of proteasome-active sites. Cell Biochem Biophys, 60, 77–88.

45. Kina, E., Larouche, J.D., Thibault, P. and Perreault, C. (2025) The cryptic immunopeptidome in health and disease. Trends Genet, 41, 162–169.

46. Navarro, A.P. and Cheeseman, I.M. (2022) Identification of a Golgi-localized peptide reveals a minimal Golgi-targeting motif. 33, ar110.

47. Brownfield, B.A. and Fromme, J.C. (2025) Structural insights into traffic through the Golgi complex. Current Opinion in Cell Biology, 94, 102505.

48. de Breyne, S., Monney, R.S. and Curran, J. (2004) Proteolytic Processing and Translation Initiation: TWO INDEPENDENT MECHANISMS FOR THE EXPRESSION OF THE SENDAI VIRUS Y PROTEINS. Journal of Biological Chemistry, 279, 16571–16580.

49. Hettmann, C., Herm, A., Geiter, A., Frank, B., Schwarz, E., Soldati, T. and Soldati, D. (2000) A dibasic motif in the tail of a class XIV apicomplexan myosin is an essential determinant of plasma membrane localization. Mol Biol Cell, 11, 1385–1400.

50. Legrand, N., Jaquier-Gubler, P. and Curran, J. (2015) The impact of the phosphomimetic eIF2αS/D on global translation, reinitiation and the integrated stress response is attenuated in N2a cells. Nucleic Acids Res, 43, 8392–8404.

51. Kisselev, A.F. (2021) Site-Specific Proteasome Inhibitors. Biomolecules, 12.

52. Käll, L., Storey, J.D. and Noble, W.S. (2008) Non-parametric estimation of posterior error probabilities associated with peptides identified by tandem mass spectrometry. Bioinformatics, 24, i42–i48.

53. Nesvizhskii, A.I., Keller, A., Kolker, E. and Aebersold, R. (2003) A statistical model for identifying proteins by tandem mass spectrometry. Anal Chem, 75, 4646–4658.

