## Supplemental Results for "The microprotein SEP^53BP1^: its bizarre mode of translational expression and intracellular behaviour"

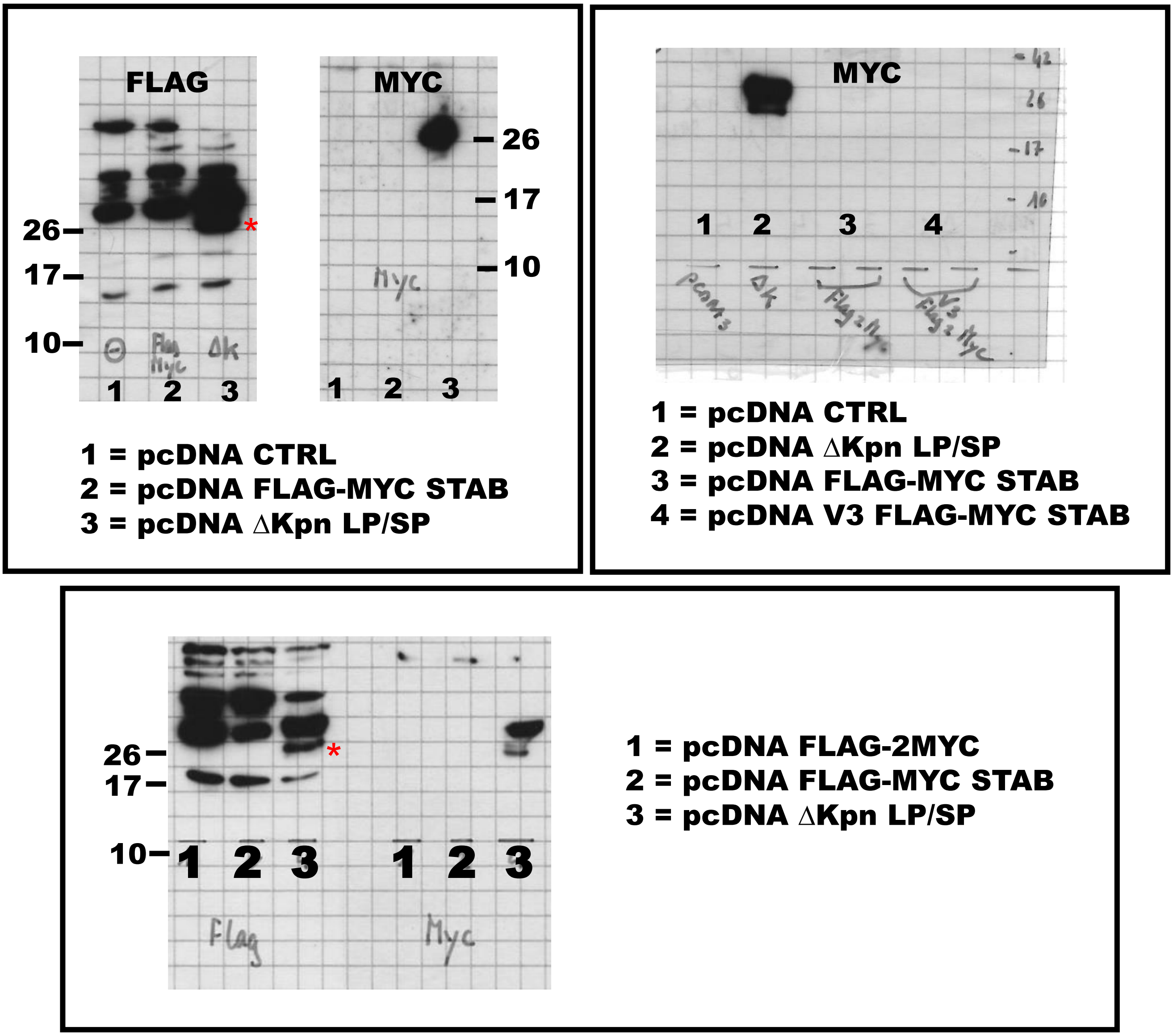


**Figure S1: Attempts to transiently express synthetic microproteins.** The figure includes a series of panels in which we attempted to detect synthetic microproteins carrying FLAG and MYC tags in transfected HEK293T cells using immunoblots with anti-MYC and anti-FLAG Abs. The positive control was supplied by a FLAG/MYC construct, ΔKpn LP/SP, as described in (31). The red star indicates the position of the control signal on the anti-FLAG immunoblots.

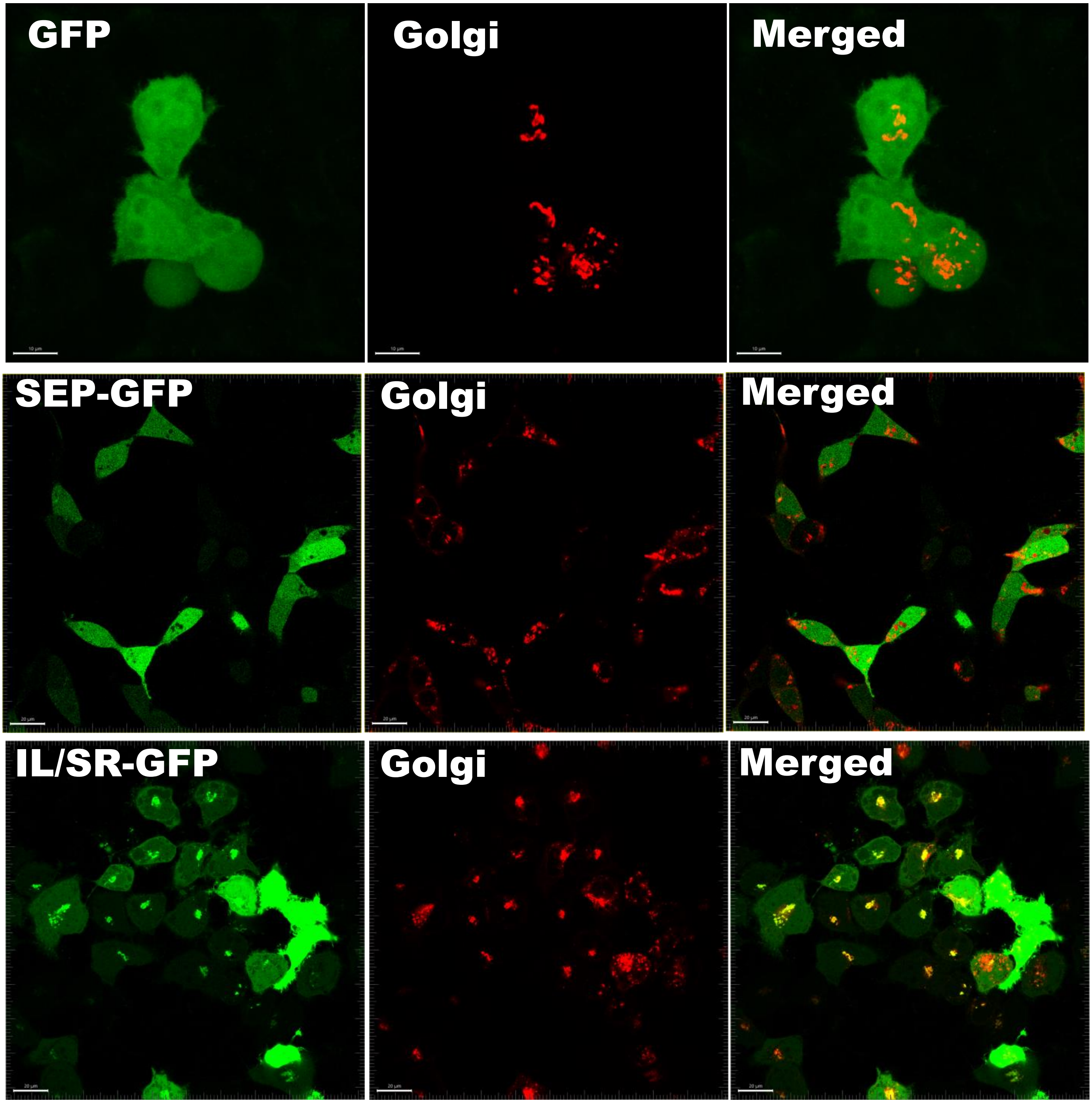


**Figure S2: Live cell images and co-localisation with the Golgi.** Cells were transfected with pmCherry-N1-GalT (which expresses the Golgi marker GalT-mCherry, GalT encoding the amino acids 1-81 from β1,4-galactosyltransferase) and the pcDNA GFP plasmids as indicated in each panel. Live cell images were recorded at 20 hrs post-transfection and analysed using the Imaris software.

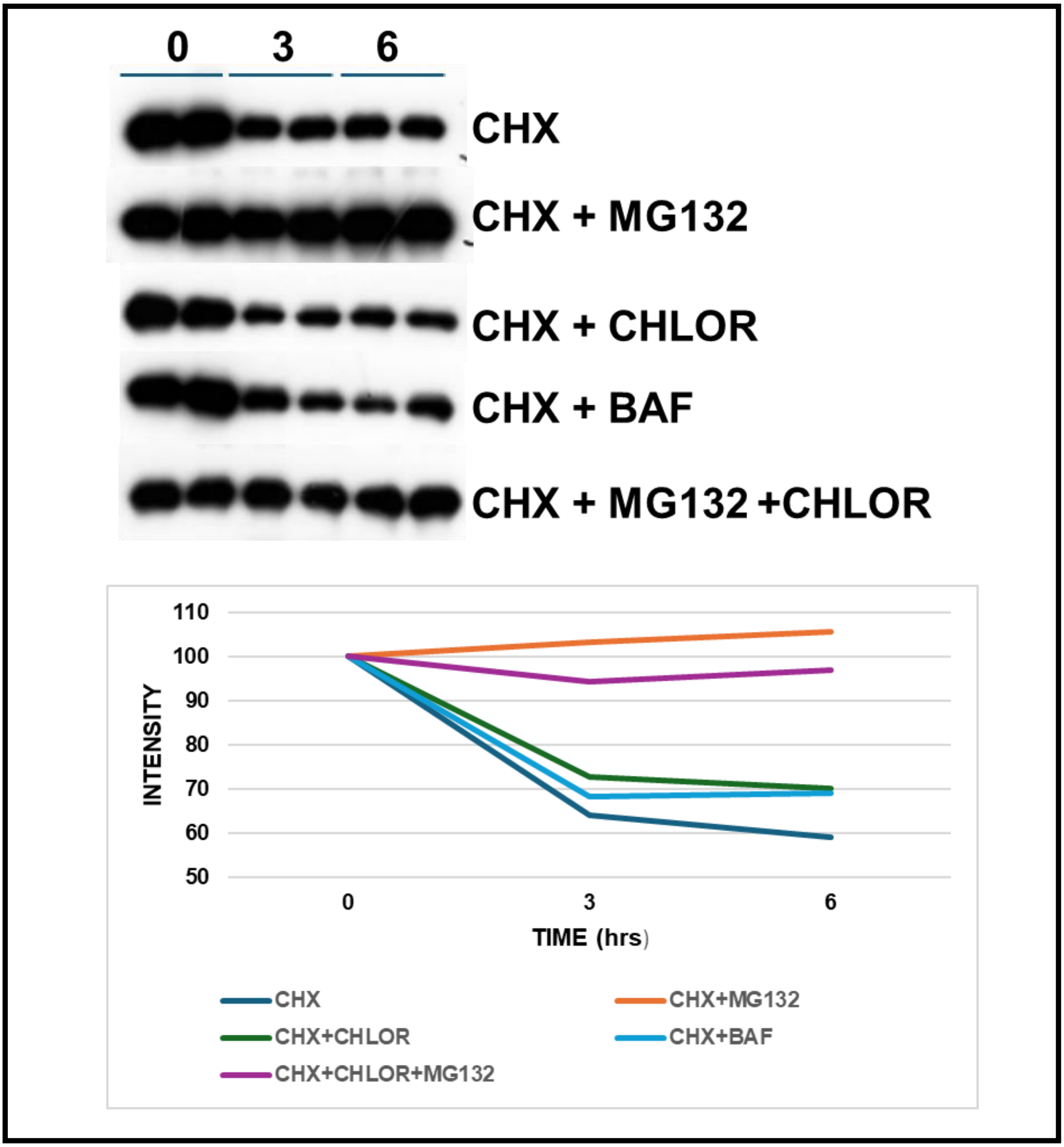


**Figure S3: Half-life studies on the GFP-SEP fusion protein.** (A). HEK293T cells transfected with a pcDNA3 construct expressing GFP-SEP were treated with cycloheximide (CHX: 100 μM) with or without the drugs as listed (MG132 10 μM: CHLOR = chloroquine 100 μM : BAF = bafilomycin 100 nM) at 20 hrs post-transfection. Cells (duplicates) were subsequently lysed at the times (hrs) indicated. Protein expression was monitored by immunoblotting with an anti-GFP Ab (upper panel). Bands were quantitated using ImageJ and the mean value was plotted (lower panel).

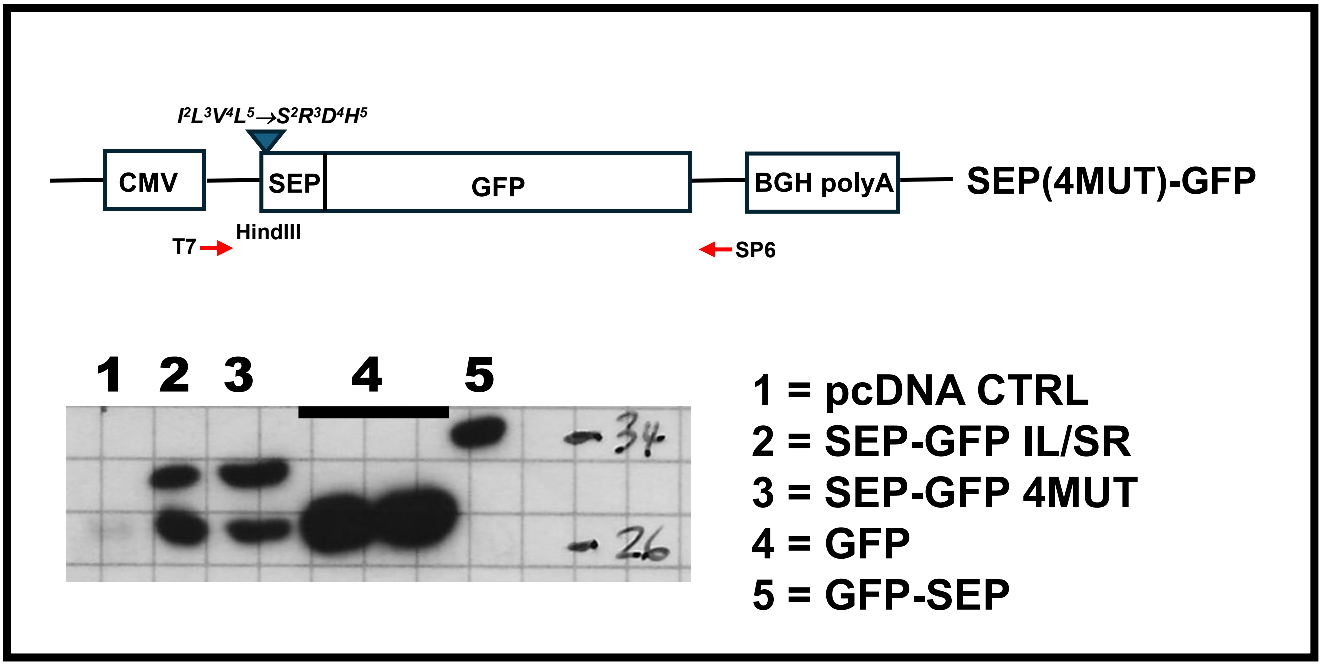


**Figure S4: Expression of the SEP-GFP N-terminal 4 MUT construct.** The upper panel is a schematic representation of the 4 MUT SEP-GFP construct in which the non-polar amino acids 2 to 5 of SEP were changed to polar charged. This construct plus the others indicated were transiently expressed in HEK293T cells. The expression pattern (lower panel) was analysed by immunoblotting using an anti-GFP Ab.
